# Supporting global biodiversity assessment through high-resolution macroecological modelling: Methodological underpinnings of the BILBI framework

**DOI:** 10.1101/309377

**Authors:** Andrew J Hoskins, Thomas D Harwood, Chris Ware, Kristen J Williams, Justin J Perry, Noboru Ota, Jim R Croft, David K Yeates, Walter Jetz, Maciej Golebiewski, Andy Purvis, Tim Robertson, Simon Ferrier

## Abstract

**Aim:** Global indicators of change in the state of terrestrial biodiversity are often derived by intersecting observed or projected changes in the distribution of habitat transformation, or of protected areas, with underlying patterns in the distribution of biodiversity. However the two main sources of data used to account for biodiversity patterns in such assessments – i.e. ecoregional boundaries, and vertebrate species ranges – are typically delineated at a much coarser resolution than the spatial grain of key ecological processes shaping both land-use and biological distributions at landscape scale. Species distribution modelling provides one widely used means of refining the resolution of mapped species distributions, but is limited to a subset of species which is biased both taxonomically and geographically, with some regions of the world lacking adequate data to generate reliable models even for better-known biological groups.

**Innovation:** Macroecological modelling of collective properties of biodiversity (e.g. alpha and beta diversity) as a correlative function of environmental predictors offers an alternative, yet highly complementary, approach to refining the spatial resolution with which patterns in the distribution of biodiversity can be mapped across our planet. Here we introduce a new capability – BILBI (the Biogeographic Infrastructure for Large-scaled Biodiversity Indicators) – which has implemented this approach by integrating advances in macroecological modelling, biodiversity informatics, remote sensing and high-performance computing to assess spatial-temporal change in biodiversity at ~1km grid resolution across the entire terrestrial surface of the planet. The initial implementation of this infrastructure focuses on modelling beta-diversity patterns using a novel extension of generalised dissimilarity modelling (GDM) designed to extract maximum value from sparsely and unevenly distributed occurrence records for over 400,000 species of plants, invertebrates and vertebrates.

**Main conclusions:** Models generated by BILBI greatly refine the mapping of beta-diversity patterns relative to more traditional biodiversity surrogates such as ecoregions. This capability is already proving of considerable value in informing global biodiversity assessment through: 1) generation of indicators of past-to-present change in biodiversity based on observed changes in habitat condition and protected-area coverage; and 2) projection of potential future change in biodiversity as a consequence of alternative scenarios of global change in drivers and policy options.

## INTRODUCTION

Continued growth in human populations around the world is intensifying demands on our natural environment. Coupled with the effects of anthropogenic climate change, the potential for large-scale modification and loss of our planet’s remaining biological diversity seems ever more likely (Pereira *et al.*, 2010). To combat this ongoing decline, governments have agreed to multi-lateral policy goals which aim to limit, reduce or halt biodiversity loss and environmental degradation. The Convention on Biological Diversity (CBD) Strategic Plan for Biodiversity 2011-2020 and the associated Aichi Biodiversity Targets are one such policy framework that sets near-future targets across five strategic goals addressing ultimate drivers of biodiversity loss, proximate pressures, management responses, benefits to people, and implementation challenges (CBD, 2010). More recently the Sustainable Development Goals (SDGs) adopted by the United Nations promote a healthy and sustainable future both for humans and for our environment, including all “life on land” and “life below water” (UN, 2015), while the latest multi-lateral agreement to limit anthropogenic climate change, ratified in Paris in 2015, includes statements to limit the loss of natural habitat (through deforestation) with indirect consequences for biodiversity (Citroen *et al.*, 2016).

Efficient planning of actions to achieve biodiversity-related goals and targets under these policy processes, and effective tracking of progress towards this achievement, requires the ability to measure the present state of biodiversity, detect trends of recent change, and project the potential future state of biodiversity expected under alternative policy options in a globally consistent way. Unfortunately our ability to report or project indicators of change for many aspects of biodiversity is still limited by an inability to observe or infer changes in ecological communities directly from currently available global datasets (Ferrier, 2011). Indicators of change employed in biodiversity assessments are most often derived by intersecting observed or projected changes in the distribution of habitat loss and degradation, or of protected areas, with underlying patterns in the distribution of biodiversity (e.g. Tittensor *et al.*, 2014; Butchart *et al.*, 2015)

Two sources of global data on terrestrial biodiversity patterns have been used most commonly in the derivation of protected-area and habitat indicators. The first of these is the World Wildlife Fund’s mapping of 867 terrestrial ecoregions, defined as “relatively large units of land containing a distinct assemblage of natural communities and species, with boundaries that approximate the original extent of natural communities prior to major land-use change” (Olson *et al.*, 2001). Ecoregions have long provided a convenient and well-respected foundation for assessing changing patterns of protected-area coverage and habitat transformation around the world (e.g. Watson *et al.*, 2016). However, as indicated by the above definition, ecoregions are typically delineated at a much coarser resolution than the spatial grain of key ecological processes shaping both land-use and biological distributions at the landscape scale (Londoño-Murcia *et al.*, 2010; Calderón-Patrón *et al.*, 2016; Serrano *et al.*, 2018).

Using ecoregions as fundamental spatial units for assessing impacts of protected-area coverage and habitat transformation on biodiversity assumes that all biological elements (e.g. species) within an ecoregion will be equally affected by these activities. Yet, in reality, fine-scaled spatial heterogeneity in abiotic environmental attributes (e.g. terrain, soils, climate) within an ecoregion will tend to bias human uses to particular parts of the region (e.g. a greater likelihood of agriculture in flatter, more fertile environments) (Fig. 1a). Since these same environmental attributes shape natural distributions of species at landscape scale (Fig. 1b), impacts of any given land-use change within an ecoregion will tend to be biased towards a subset of the species occurring within that region (Fig. 1c). This means that protected-area or habitat indicators derived using ecoregions as the fundamental units of analysis risk under-or over-estimating implications of protection or habitat transformation for biodiversity contained within these regions (Ferrier *et al.*, 2004; Londoño-Murcia *et al.*, 2010).

**Figure 1:**
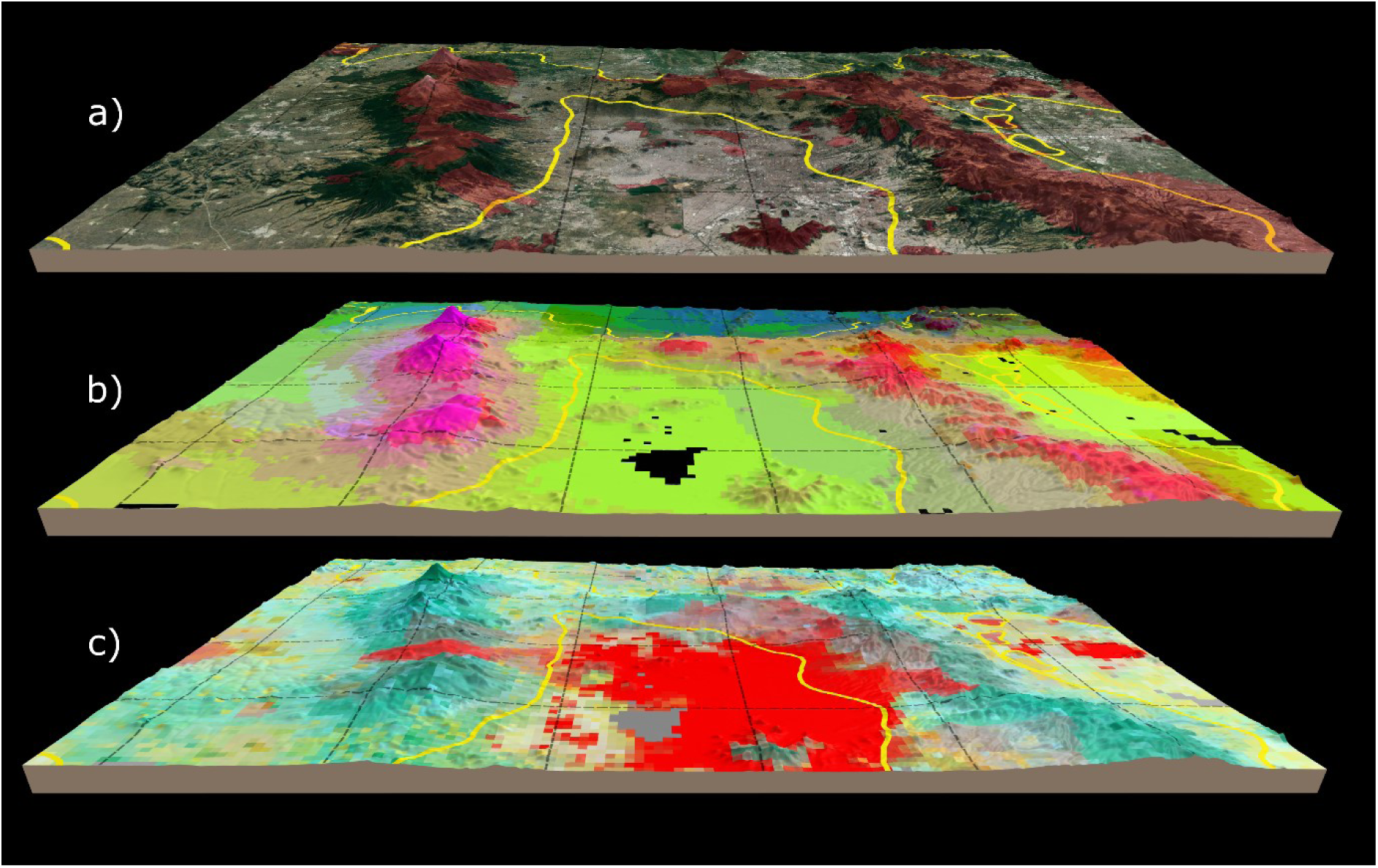
Depiction of the problem of scale when making biodiversity assessments. (a) shows a true colour satellite overlay of the region surrounding Mexico city with typical resolutions for global analyses – a 30 km^2^ grid (black) and ecoregional boundaries (yellow) – and protected areas overlayed in red. Note the geographic/topographic biases in placement of both protected areas and urban development. (b) shows the same region but the different ecological communities (defined by similarity between communities in each 1 km^2^ pixel) where similar colours represent similar communities. (c) shows land-use from Hoskins *et al.* (2016) where red depicts urbanisation, yellow represents cropping regions and green shows natural environments – note: colours show proportional values of land-use for each pixel with the transparency set as the proportion of each land-use per cell, as such, blended colours represent pixels with mixed land-uses.

The other major source of global data on biodiversity patterns commonly used for deriving indicators – i.e. extent-of-occurrence range maps for terrestrial vertebrate species (e.g. Jenkins *et al.*, 2013) – presents similar spatial-resolution challenges. As for ecoregions, this data source has, over recent years, enabled the derivation of a wide variety of indicators, and has also underpinned numerous macroecological analyses of global biodiversity patterns. However the relatively coarse resolution of most range maps, and the reality that species occupy only those parts of their overall range offering suitable environmental conditions, has led some workers to suggest that these data should not be employed at a grid resolution finer than 1 degree, or approximately 100km x 100km near the equator (Hurlbert & Jetz, 2007). This again is a resolution far coarser than the spatial grain of key ecological processes shaping land-use and biological distributions at landscape scale.

Species distribution modelling (SDM) provides one widely used means of refining the resolution of mapped species distributions, by using fine-resolution environmental surfaces to characterise and spatially project a species’ niche space (Elith & Leathwick, 2009). This can be achieved either by using known occurrence records to fit a correlative model predicting occurrence of a given species as a mathematical function of multiple environmental variables, or through deductive modelling in which occurrence is predicted using simple rule-based descriptions of environmental suitability derived from expert knowledge (Ferrier, 2002). Distributions predicted using SDM can be used either directly in assessments, or combined with mapped species ranges, where available (e.g. for vertebrates), thereby providing refined mapping of the expected distribution of each species within its known range (Merow *et al.*, 2017). However, regardless of the precise SDM technique employed, application of this general approach is restricted to species for which either there is a sufficient number of occurrence records available to develop a correlative model, or there is sufficient expert knowledge of the species’ habitat requirements to develop a deductive model. This capacity is therefore limited to a subset of species which is biased both taxonomically and geographically, with some regions of the world lacking adequate data to generate reliable SDMs even for better-known biological groups such as vertebrates, let alone for invertebrates and plants (Meyer *et al.*, 2015).

Here we adopt an alternative, yet highly complementary, approach to integrating species-occurrence records with fine-scaled environmental surfaces. This allows us to refine the spatial resolution with which patterns in the distribution of biodiversity can be mapped across our planet. Rather than attempting to model distributions of individual species, this approach instead focuses on modelling, and thereby mapping, collective properties of biodiversity as a correlative function of environmental predictors. Macroecological modelling of spatial variation in alpha diversity, particularly of variation in local species richness, has a relatively long history of application in ecology and conservation biology (e.g. Francis & Currie, 1998). However, with increasing awareness that the total (gamma) diversity encompassed by any set of areas (e.g. in a conservation reserve system) will typically depend more on the extent to which these areas complement one another in terms of species composition, than it does on the richness of individual areas, macroecological modelling is now placing greater emphasis on modelling patterns of beta diversity in addition to those of alpha diversity (Ferrier & Guisan, 2006; D’Amen *et al.*, 2017).

We here introduce a new capability for global biodiversity assessment – BILBI (the Biogeographic Infrastructure for Large-scaled Biodiversity Indicators) – underpinned by macroecological modelling of collective properties of biodiversity. The initial implementation of this infrastructure relies strongly on modelling of beta diversity patterns using an extension of one particular technique – generalised dissimilarity modelling (GDM; Ferrier *et al.*, 2007) – applied to readily available biological and environmental datasets. The overall framework is, however, designed to be sufficiently generic and flexible to allow incremental refinement and addition of modelling techniques and datasets into the future. This capability is also intended to complement, rather than compete with, other approaches to global biodiversity assessment, including those focussed on individual species (e.g. Jetz *et al.*, 2012). Species-level approaches will always play a vital role in biodiversity assessment for better-known biological groups, and especially for species of particular conservation concern within these groups. However the approach described here has potential to add significant value to such species-based assessments by: 1) allowing more effective use of data for highly-diverse biological groups, containing large numbers of species but with few records per species; and 2) enabling robust extrapolation of expected patterns across poorly-sampled regions, even where the particular species occurring in these regions are unknown or unsurveyed.

## GENERAL FRAMEWORK

The BILBI modelling framework (Fig. 2) integrates advances in macroecological modelling, biodiversity informatics, remote sensing and high-performance computing to assess spatio-temporal change in biodiversity at 30-arcsecond (approximately 1km) grid resolution across the entire terrestrial surface of the planet, excluding Antarctica (above 60∼S). Best-available data on observed occurrences of species within defined biological groups (e.g. all vascular plants) are used to fit correlative models describing patterns in the distribution of biodiversity as a function of fine-scaled spatial variation in climate, terrain and soils, within major habitat types (biomes) and biogeographic realms. These patterns are mapped as spatially-complete gridded surfaces by interpolating and, where necessary, extrapolating predictions from the fitted models. The resulting surfaces describe patterns in the spatial distribution of biodiversity which would be expected in the absence of anthropogenic habitat transformation. These modelled patterns then serve as the foundation for two subsequent pathways of analysis in the BILBI framework (Fig. 2).

**Figure 2:**
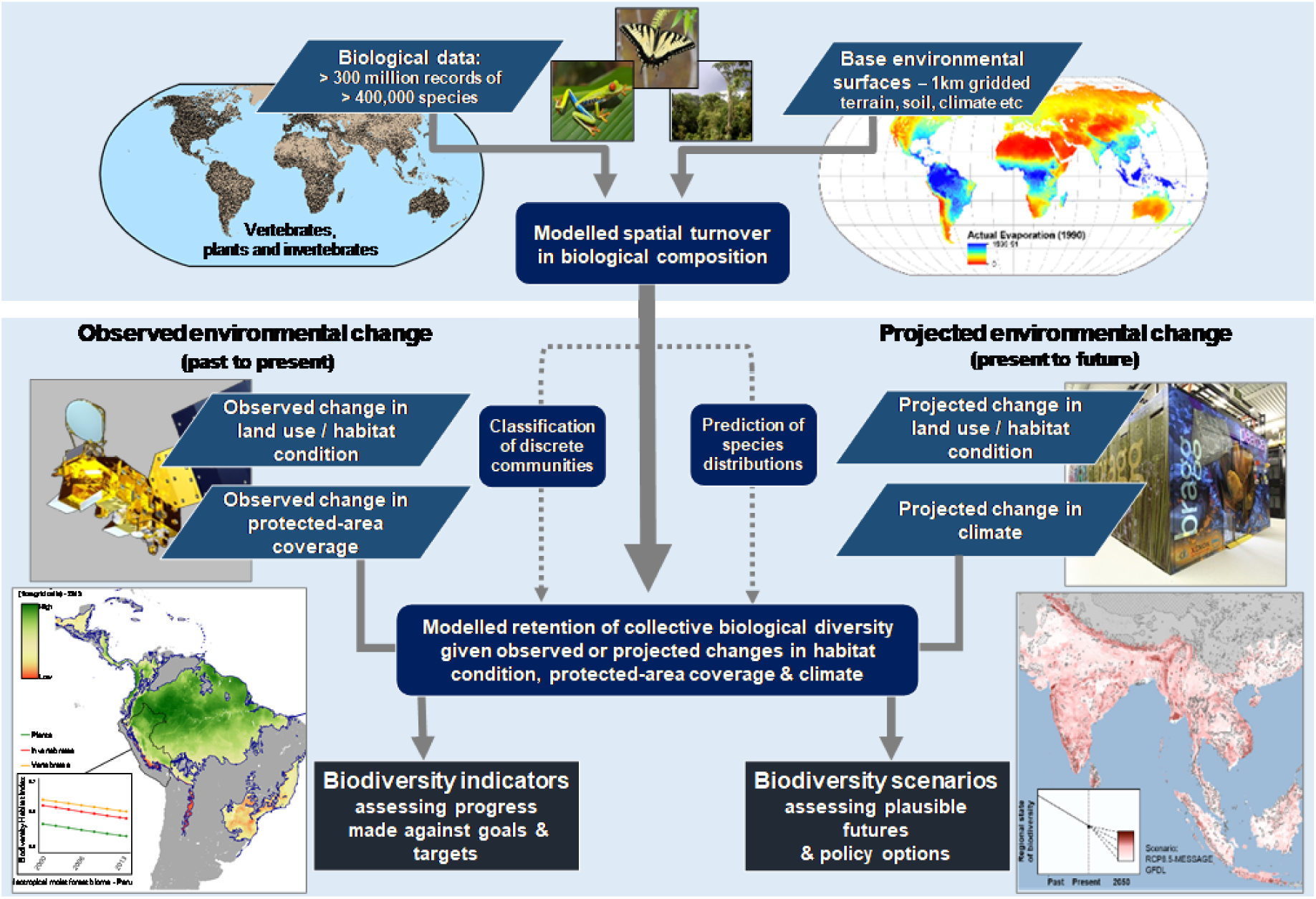
The basic structure of the BILBI modelling framework, showing how our initial modelling of compositional turnover can follow to separate analysis pathways to produce either indicators of recent change in community composition or possible future scenarios of change to composition.

In the first pathway these patterns of biodiversity distribution are overlayed with observed changes in pressures (direct drivers) – particularly changes in habitat condition resulting from land-use change – or in management responses, such as the establishment of protected areas, to generate indicators of biodiversity change (e.g. for reporting progress towards the CBD’s Aichi Targets). In the second pathway, observed changes are replaced by projected changes in pressures and responses into the future. This enables application of BILBI in translating alternative scenarios of global change, and associated policy or management options, into expected consequences for the future persistence of biodiversity. In assessing such scenarios the BILBI framework allows consideration both of impacts mediated by changes in habitat condition, resulting for example from projected land-use change, and of potential impacts of climate change on community composition. The latter is predicted through space-for-time substitution of climate covariates in BILBI’s correlative models of spatial biodiversity distribution.

In the remainder of this paper we describe our initial implementation of the BILBI framework, focusing primarily on the foundational modelling of spatial patterns in the global distribution of terrestrial biodiversity. The two analytical pathways flowing from this foundation – relating to indicator generation, and scenario analysis respectively (Fig. 2) – will be addressed in detail in subsequent papers. As alluded to above, our modelling of biodiversity patterns in the initial implementation of BILBI has focused on describing and predicting spatial turnover in species composition – i.e. patterns of beta diversity. However, our longer-term intent is to extend this approach to accommodate joint modelling of both alpha and beta diversity; and to integrate next-generation techniques for achieving this as they become operational.

## INITIAL GLOBAL IMPLEMENTATION

### Modelling compositional turnover using presence-only data

Generalised dissimilarity modelling is a nonlinear regression technique for modelling the turnover in species composition between two sites as a function of environmental differences between, and geographical separation of, these sites. This technique accommodates two types of nonlinearity commonly encountered in large-scaled analyses of compositional turnover. The curvilinear relationship between increasing environmental or geographical distance, and observed compositional dissimilarity, between sites is addressed through the use of appropriate link functions in a generalised linear modelling framework. Variation in the rate of compositional turnover at different positions along environmental gradients is addressed by transforming these gradients using smooth monotonic functions fitted to the training data (Ferrier *et al.*, 2007). The response variable in a standard GDM model is typically a measure of between-site compositional dissimilarity, calculated from lists of species observed at each of the two sites, using indices such as Sørensen or βsim (e.g. Jones *et al.*, 2013; König *et al.*, 2017). However, one of the biggest challenges in applying GDM globally has been that a large proportion of available species-occurrence data are presence-only rather than presence-absence in nature. Most occurrence records accessible through major data infrastructures, such as the Global Biodiversity Information Facility (GBIF), have been generated through geo-referencing of specimens from natural-history collections, or from relatively opportunistic field observations of individual species, rather than from planned inventories systematically recording all species present at a given site (Isaac & Pocock, 2015). Such data are not well suited to estimating compositional dissimilarity between sites, particularly in areas with lower sampling effort. This is because estimates of compositional dissimilarity will be inflated, to a varying yet unknown extent, by false absences of species at each of the sites concerned (Beck *et al.*, 2013).

In implementing the BILBI framework we have addressed this problem by modifying GDM to work with a binary response variable, defined in terms of matches versus mismatches in species identity, for pairs of individual species observations (where a “species observation” is the recorded presence of a particular species at a particular site). The probability that a species randomly drawn from site *i* has the same identity as a species randomly drawn from site *j* is expected to be a function of both the total number of species actually occurring at each of the two sites (alpha diversity) and the number of species at each site which are unique to that site, because they do not occur at the other site (beta diversity), following the expression:

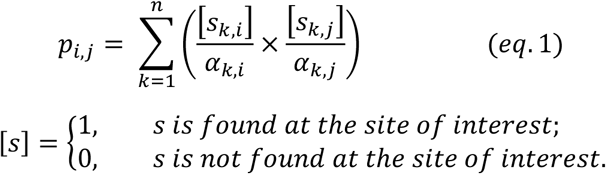

where *s* is the identity of an individual species belonging to the combined list of *n* species occurring at either one, or both, of the sites and α is the number of species found at a particular site. As the quantity being summed reduces to 0 when a given species is not shared by both sites, this equation can be simplified to *p*_*i,j*_ = (1/[α_*i*_ α_*j*_])*c*, where *c* is the number of species shared between the two sites.

Using this understanding, we fit a modified form of GDM in which the standard response variable, describing the compositional dissimilarity between two sites (on a continuous scale between 0 and 1), is now replaced by a binary response variable describing the match (0) or mismatch (1) in species identity of a randomly drawn pair of species observations from the two sites. The probability of a mismatch in species identity is the complement of *pi,j* from above – i.e. 1-*p*_*i,j*_. This probability is modelled as a non-linear function of environmental covariates (predictors), in a similar manner to that employed in standard GDM model-fitting (Ferrier *et al.*, 2007). However, due to the binary nature of the response when working with pairs of species observations, the negative exponential link function traditionally used in GDM is replaced by the logit link function. The overall form of an observation-pair GDM (*obs-pair*GDM) fitted to *m* environmental covariates is therefore:

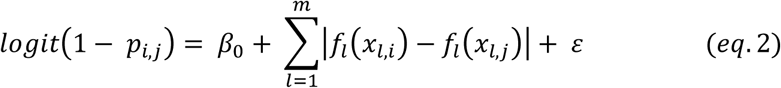

where |*f*_*l*_(*x*_*l,i*_) − *f*_*l*_(*x*_*l,j*_)| represents the separation of a pair species observations *i* and *j* along a nonlinear function of environmental covariate *l* fitted using monotonic I-splines (Ferrier *et al.*, 2007). This is not yet an estimate of the turnover in species composition between two sites. As we showed above, *p_i,j_* is a function not only of compositional turnover, but also of the richness of species at the two sites concerned. However, if we can estimate the mean species richness of these sites then we can decompose *p_i,j_* into an estimate of compositional turnover between the sites (*d*_*i,j*_) using:

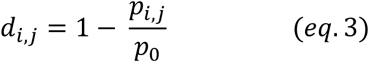

where *p*_0_ represents an estimate of the probability that a randomly drawn pair of species from a pair of identical sites (i.e. compositional turnover equals 0) are the same. To enable fitting we estimate *p*_0_ from the intercept of our model – i.e. the point where environmental separation between sites is 0 and thus the sites are treated as the same. For any subsequent analyses any prediction of *d*_*i,j*_ between a pair of sites can be treated as the expected proportion of species occurring at one site which do not also occur at the other site (averaged across the two sites), and is therefore effectively an estimate of the Sørensen index.

By fitting our models to pairs of individual species observations this method avoids the biases that can result from modelling community data where the inventory of species at sites is incomplete. We just need to satisfy the assumption that the particular species recorded as present at a given site constitute a random sample drawn from all species actually occurring at that site. In its current form the method also assumes that local species richness (alpha diversity) remains reasonably constant across the region of interest – i.e. that the number of species actually occurring at individual sites (1km cells in this study) does not vary markedly across the region – and therefore that the effect of alpha diversity on the response being modelled is accounted for by the model’s intercept. As we describe in the next section, we have taken considerable care to minimise violation of this assumption by fitting separate models for different biomes and biogeographic realms. Our team is also currently developing an extension of the above approach which relaxes this assumption, and thereby explicitly models *p_i,j_* as a function of variation in both alpha and beta diversity. Preliminary testing suggests that that this approach holds considerable promise as a means of simultaneously modelling patterns of both species richness and compositional turnover from presence-only data.

### Biological inputs

Compositional-turnover models covering the terrestrial surface of the planet (above 60∼S) were developed for three biological groups – vascular plants, invertebrates and vertebrates. Species-occurrence data were obtained by first downloading all occurrence records accessible through GBIF for vascular plants, invertebrates and reptiles (GBIF.org, 20 March 2014), and all records accessible through the Map of Life (MoL; Jetz *et al.*, 2012) for birds, amphibians and mammals (as of May, 2014). All data were filtered to remove: records without accepted genus and species names (using the GBIF Backbone Taxonomy; https://www.gbif.org/dataset/d7dddbf4-2cf0-4f39-9b2a-bb099caae36c); records with a specified spatial precision greater than 10km; and records falling outside of our land mask (i.e. in the ocean or in other large water bodies). Records falling ≤ 1 km from the coastline, but belonging to terrestrial species, were moved to the nearest terrestrial cell.

For each of the three broad biological groups for which compositional-turnover models were derived – i.e. vascular plants, invertebrates and vertebrates – occurrence records were further filtered, and partitioned into taxonomic sub-groups, based on rules specific to each broad group. The main purpose of this sub-grouping was to ensure that pairs of species observations used in model fitting were of species likely to be recorded by the same community of practice of scientists and/or naturalists. For vertebrates, records were partitioned into four sub-groups – mammals, birds, reptiles and amphibians – meaning, for example, that a record of a mammal species could only be paired with another mammal record, and not with a record of a bird, reptile or amphibian. All species in these four vertebrate sub-groups were used in model fitting, except those which obtain resources primarily from the marine environment (e.g. procellariform seabirds and pinniped). For invertebrates, the 15 sub-groups employed each was associated with a relatively strong global community of practice (therefore ensuring a reasonably stable taxonomy and geographic spread of records) and contained species which are predominantly terrestrial, or are in a terrestrial phase of their life cycle when sampled. These invertebrate sub-groups, mostly arthropods, were: ants, bees, beetles, bugs, butterflies, centipedes, dragonflies, grasshoppers, millipedes, moths, snails, spiders, termites, true flies, and wasps (see Table 1 for details). For vascular plants, all species were placed in a single sub-group.

**Table 1:**
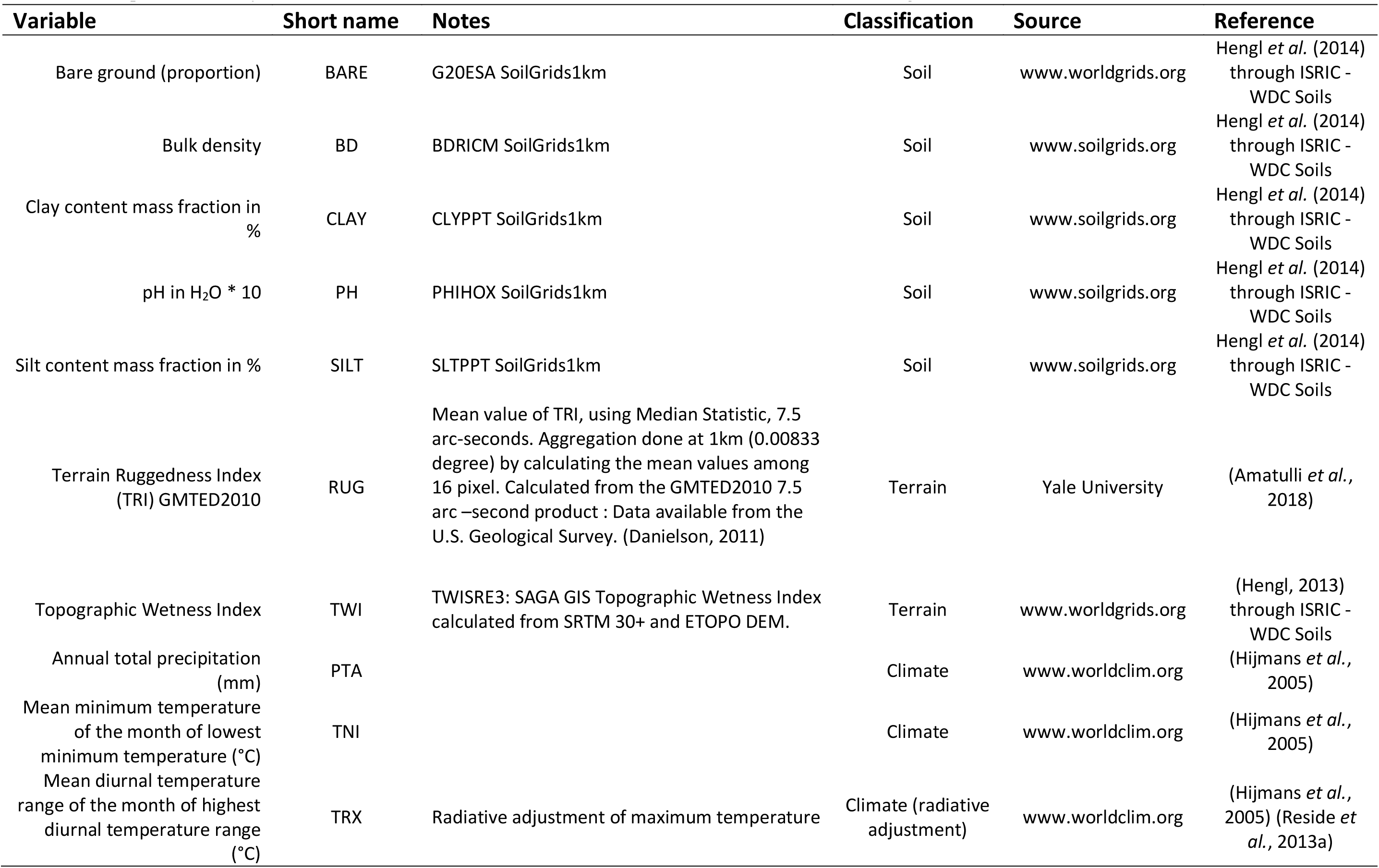
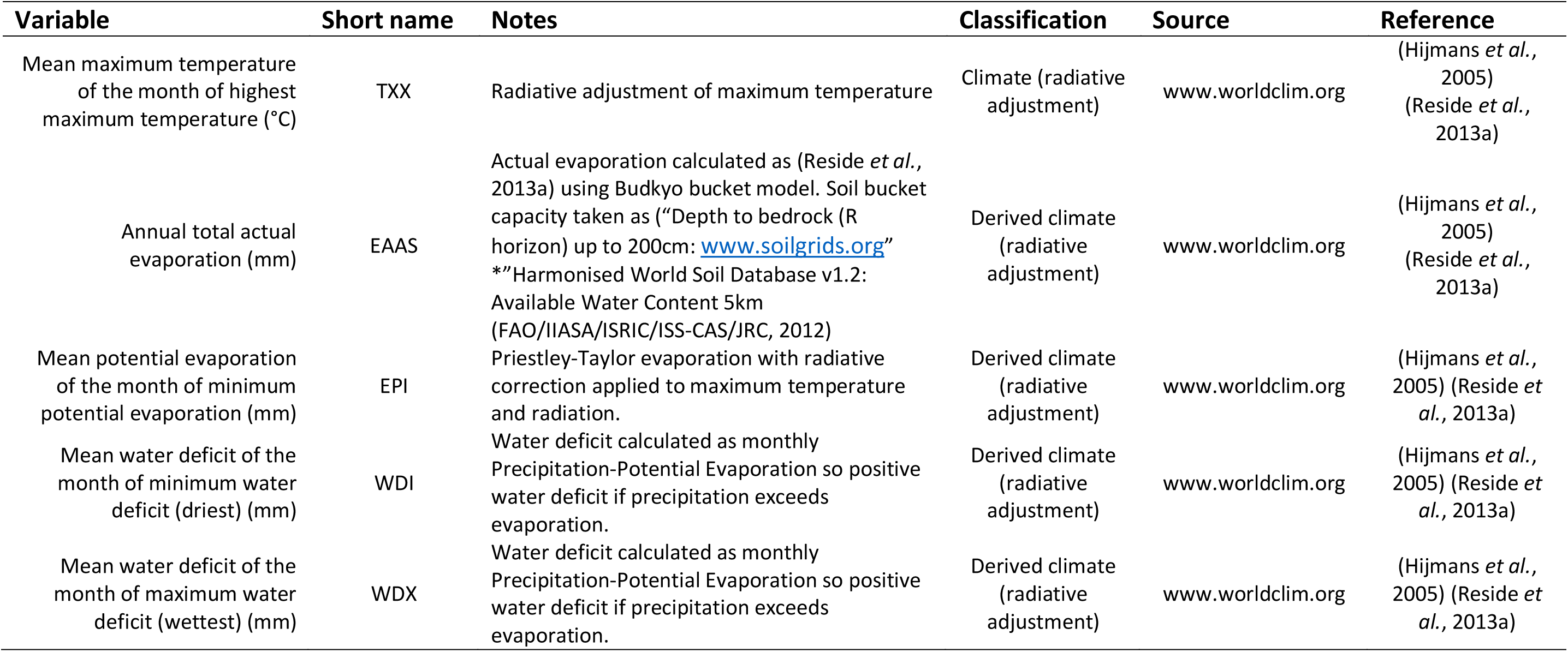
Break down of the hierarchy in observation-pair sampling for each of the three model sets used in BILBI, including the target number of observation pairs attempted for each taxonomic group and subgroup set and the number of unique species and observations available.

All species records which passed the above filtering processes were then assigned to individual 30-arcsecond resolution grid-cells concordant with the spatial grain of the environmental surfaces used in the modelling. These data were then consolidated to remove replicate species records - i.e. only a single record of any given species was retained for any given cell. A final filter was then applied to remove cells with an extremely high number of species (relative to surrounding sites), likely representing the location of biological collections (e.g. museums) rather than actual species locations. The final pool of data used for model fitting contained 106,815,923 records of a given species in a given cell, for 411,348 species (Table 1).

### Environmental inputs

Environmental covariates were selected *a priori* from the suite of possible climate, terrain and soil predictors. Individual model selection of covariate sets was not performed as this would reduce the comparability of results between models and limit our ability to generate continuous surfaces of biological composition across large areas. This *a priori* selection aimed to capture ecologically limiting factors of major importance across the world, and drew particularly on variables which had made a significant contribution to models previously fitted by our team in GDM-modelling exercises across a range of scales and biological groups. Additional criteria were that the layers provided consistent global coverage, and that they were freely available.

A standard set of 15 environmental variables were employed in all of the fitted GDM models (Table 2). This set included five soil variables (bare ground, bulk density, clay, pH, silt), two terrain variables (topographic roughness index, topographic wetness index) and eight climate variables (annual precipitation, annual minimum temperature, annual maximum temperature, maximum monthly diurnal temperature range, annual actual evaporation, potential evaporation of driest month, maximum and minimum monthly water deficit). For the climate variables it was important that these could be consistently projected into the future, and so the WorldClim elevation-adjusted data set (Hijmans *et al.*, 2005) was chosen over products incorporating remotely-sensed data. We further adjusted the temperature, evaporation and water-deficit variables for the radiative-shading effects of topography based on the GMTED2010 DEM (Danielson & Dean, 2011). Details of this adjustment, and of the associated techniques we used to derive the evaporation and water-deficit variables are provided in (Reside *et al.*, 2013b). All environmental grids were aligned to the WorldClim 30-arcsecond land-extent layer, with water bodies (defined as the Global Lakes and Wetlands Database v3 Lakes and Reservoirs: Lehner & Doll, 2004) masked out. Where necessary for the soil and terrain layers, minor information gaps were filled using a combination of extrapolation and appropriate values drawn from the literature. All methods developed and used to create these layers were designed to be applicable to both present-day climatic conditions and future climate scenarios generated from General Circulation Models (GCMs).

**Table 2:**
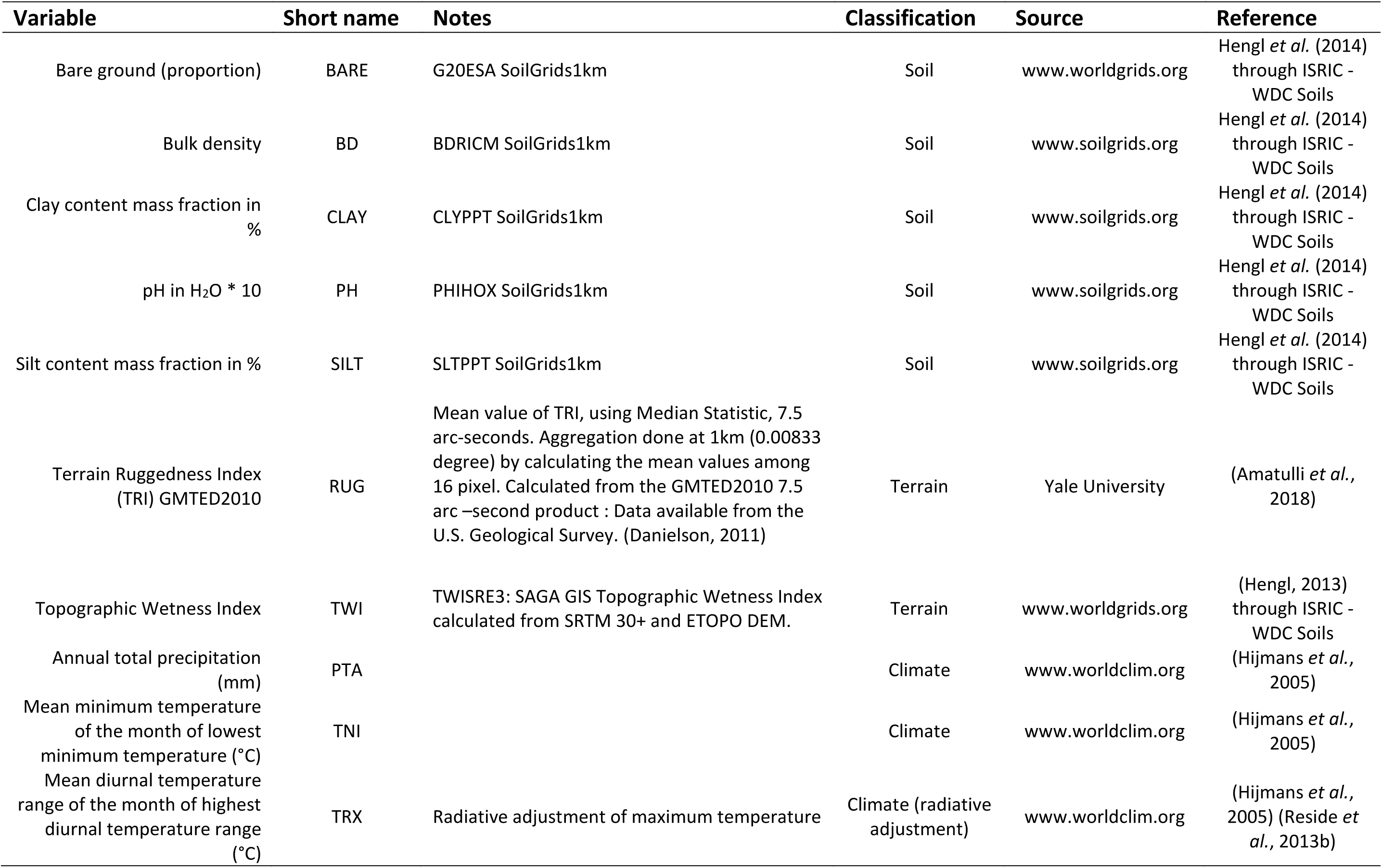
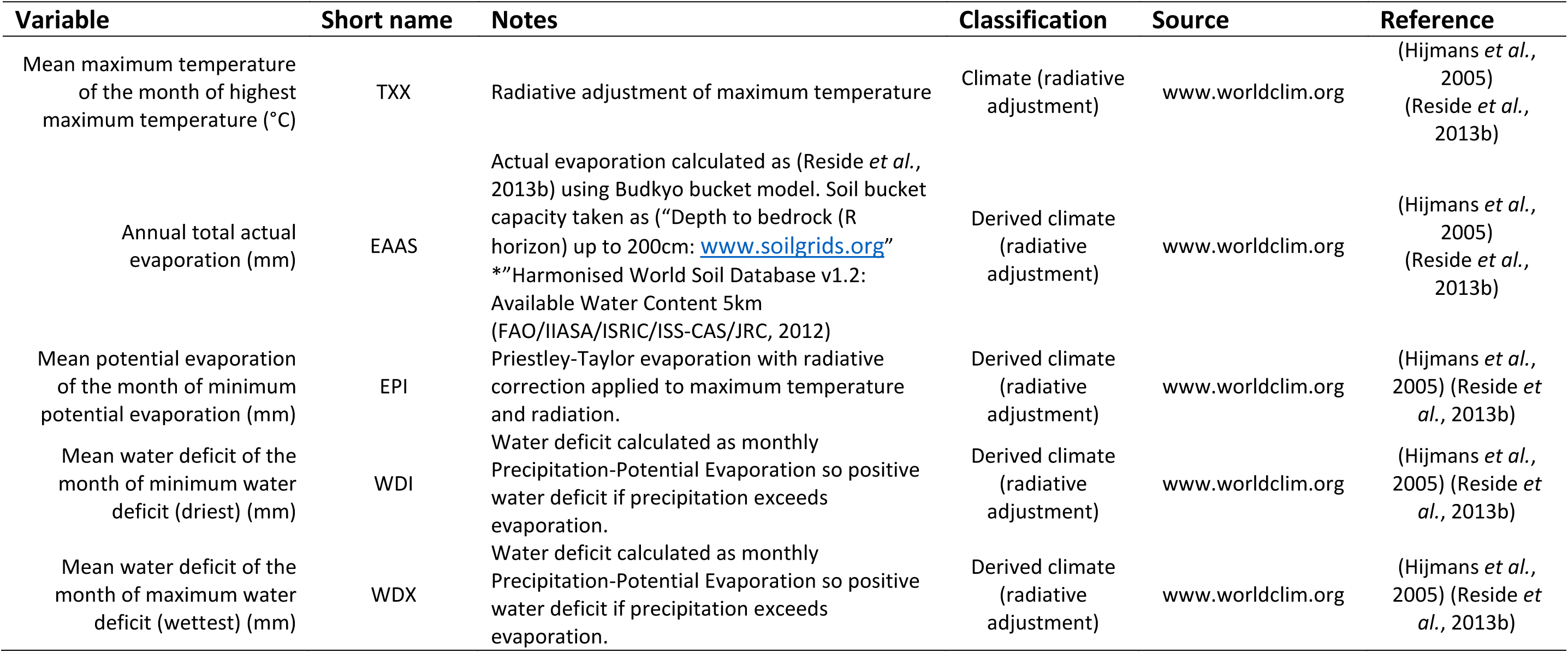
Description of data layers and their source used as environmental covariates within the BILBI modelling framework.

### Model fitting

A suite of models, describing compositional-turnover for the three biological groups (invertebrates, vascular plants and vertebrates), was generated using the *obs-pair*GDM technique (e.g. Fig. 3a-d). These models were developed within the World Wildlife Fund’s nested biogeographic-realm, biome and ecoregion framework (Olson *et al.*, 2001). A separate model was fitted for each possible pairing of the three biological groups with the 61 unique intersections of biomes and realms – referred to here as “bio-realms” – thereby yielding a total of 183 models (3 biological groups x 61 bio-realms) (Table 3).

**Table 3:**
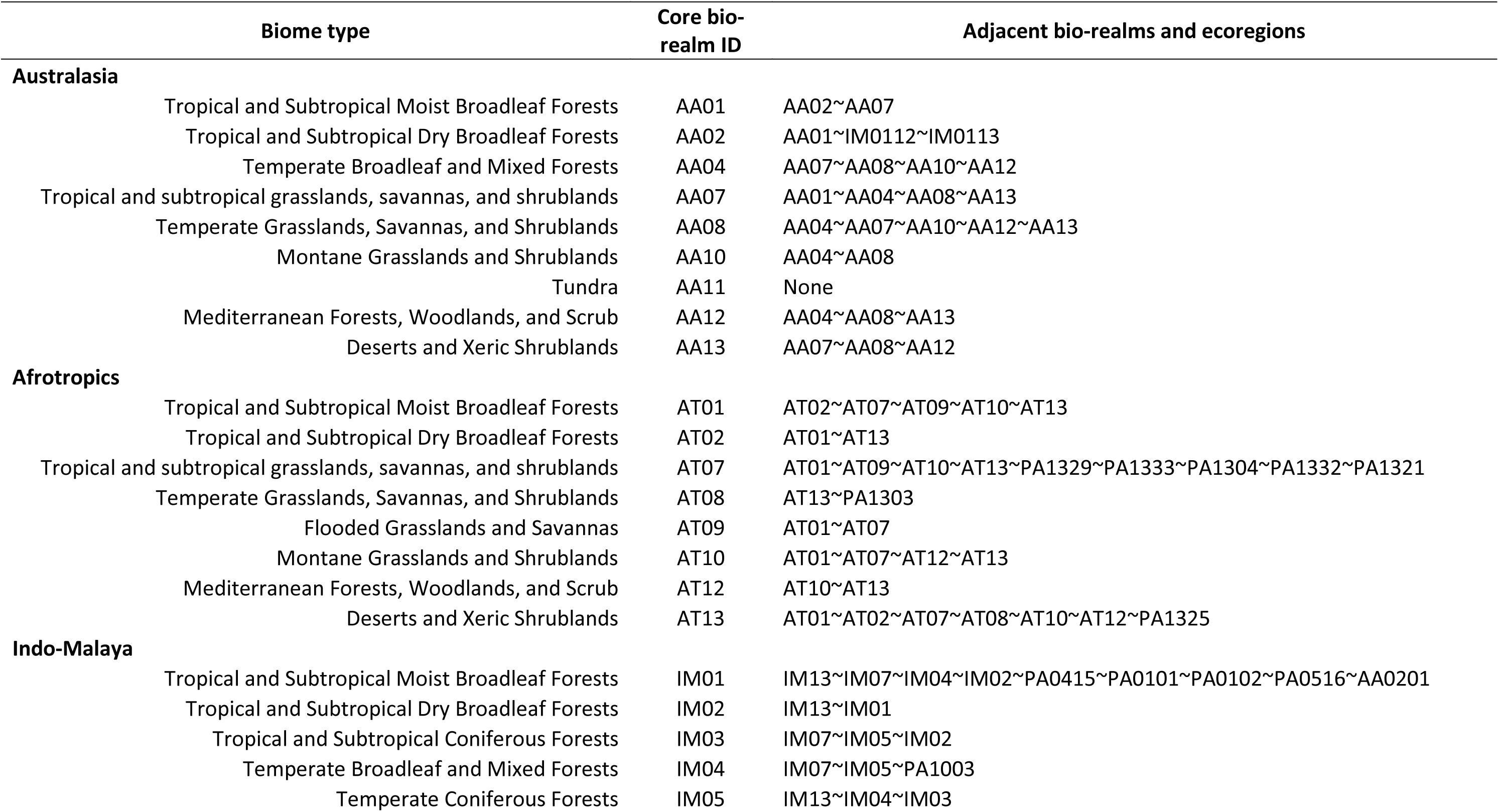
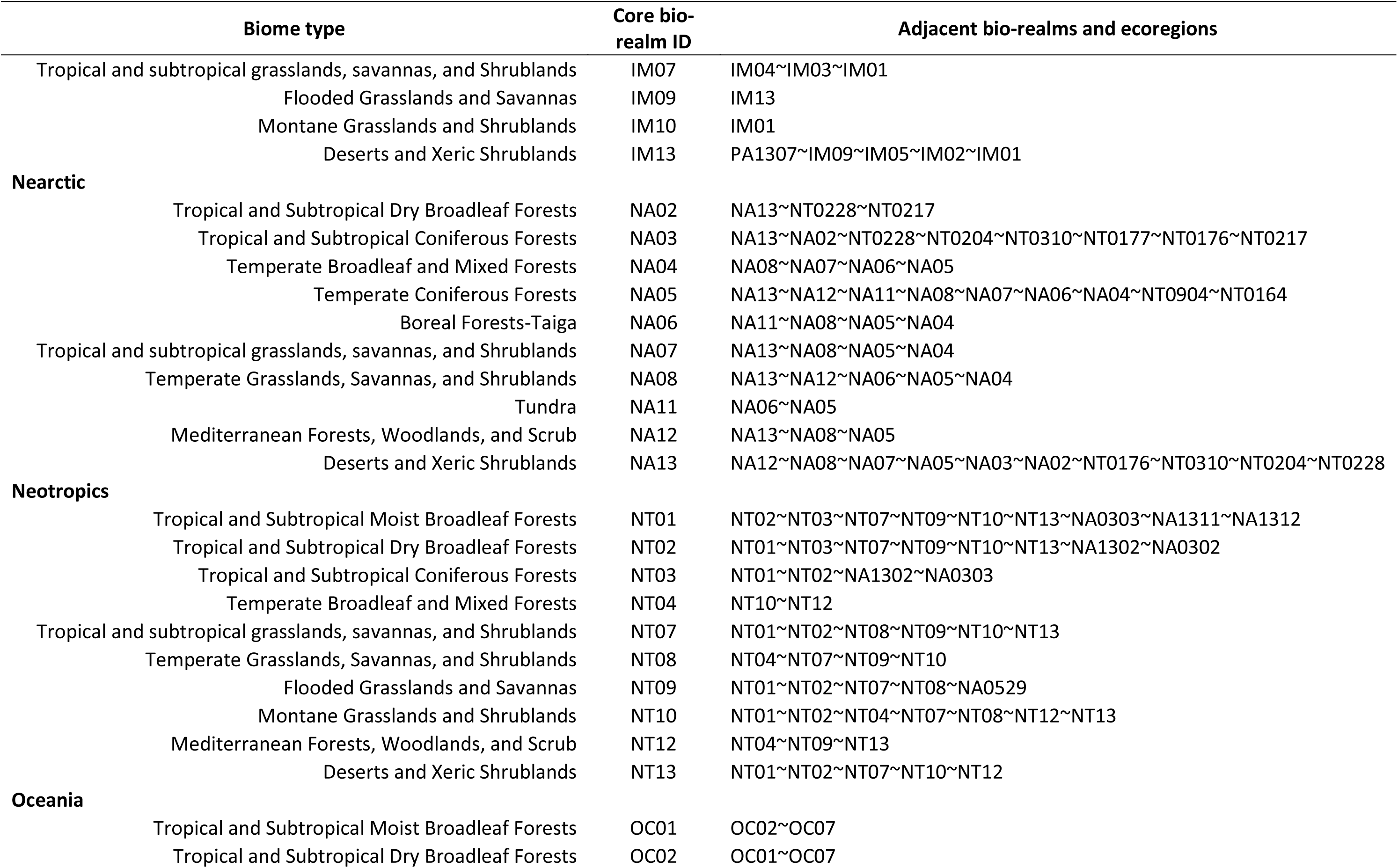
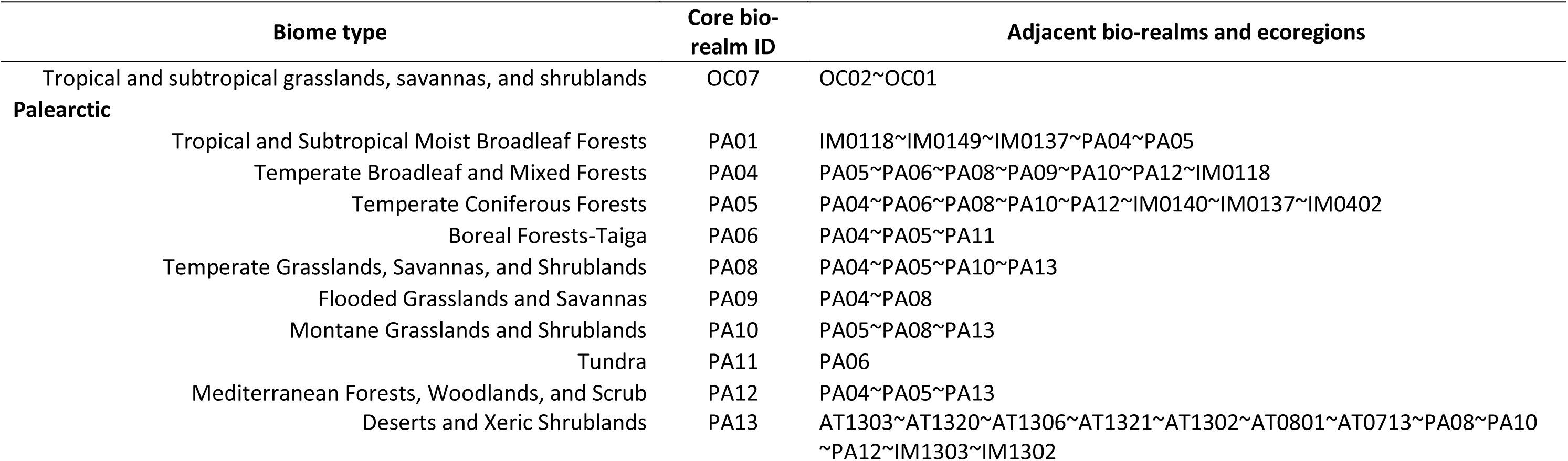
Description of the adjacent bio-realms and ecoregions used for each individual core bio-realm. Biological data were sampled with 50% of observation pairs taken from within the core region and the subsequent 50% from data encompassing both the core and adjacent regions. Bio-realm and ecoregion abbreviations follow the nomenclature established in Olson *et al.* (2001).

| Biome type | Core bio-realm ID | Adjacent bio-realms and ecoregions |
| --- | --- | --- |
| **Australasia** |  |  |
| Tropical and Subtropical Moist Broadleaf Forests | AA01 | AA02~AA07 |
| Tropical and Subtropical Dry Broadleaf Forests | AA02 | AA01~IM0112~IM0113 |
| Temperate Broadleaf and Mixed Forests | AA04 | AA07~AA08~AA10~AA12 |
| Tropical and subtropical grasslands, savannas, and shrublands | AA07 | AA01~AA04~AA08~AA13 |
| Temperate Grasslands, Savannas, and Shrublands | AA08 | AA04~AA07~AA10~AA12~AA13 |
| Montane Grasslands and Shrublands | AA10 | AA04~AA08 |
| Tundra | AA11 | None |
| Mediterranean Forests, Woodlands, and Scrub | AA12 | AA04~AA08~AA13 |
| Deserts and Xeric Shrublands | AA13 | AA07~AA08~AA12 |
| **Afrotropics** |  |  |
| Tropical and Subtropical Moist Broadleaf Forests | AT01 | AT02~AT07~AT09~AT10~AT13 |
| Tropical and Subtropical Dry Broadleaf Forests | AT02 | AT01~AT13 |
| Tropical and subtropical grasslands, savannas, and shrublands | AT07 | AT01~AT09~AT10~AT13~PA1329~PA1333~PA1304~PA1332~PA1321 |
| Temperate Grasslands, Savannas, and Shrublands | AT08 | AT13~PA1303 |
| Flooded Grasslands and Savannas | AT09 | AT01~AT07 |
| Montane Grasslands and Shrublands | AT10 | AT01~AT07~AT12~AT13 |
| Mediterranean Forests, Woodlands, and Scrub | AT12 | AT10~AT13 |
| Deserts and Xeric Shrublands | AT13 | AT01~AT02~AT07~AT08~AT10~AT12~PA1325 |
| **Indo-Malaya** |  |  |
| Tropical and Subtropical Moist Broadleaf Forests | IM01 | IM13~IM07~IM04~IM02~PA0415~PA0101~PA0102~PA0516~AA0201 |
| Tropical and Subtropical Dry Broadleaf Forests | IM02 | IM13~IM01 |
| Tropical and Subtropical Coniferous Forests | IM03 | IM07~IM05~IM02 |
| Temperate Broadleaf and Mixed Forests | IM04 | IM07~IM05~PA1003 |
| Temperate Coniferous Forests | IM05 | IM13~IM04~IM03 |

| Biome type |  | Core bio-realm ID | Adjacent bio-realms and ecoregions |
| --- | --- | --- | --- |
|  | Tropical and subtropical grasslands, savannas, and Shrublands | IM07 | IM04~IM03~IM01 |
|  | Flooded Grasslands and Savannas | IM09 | IM13 |
|  | Montane Grasslands and Shrublands | IM10 | IM01 |
|  | Deserts and Xeric Shrublands | IM13 | PA1307~IM09~IM05~IM02~IM01 |
| **Nearctic** |  |  |  |
|  | Tropical and Subtropical Dry Broadleaf Forests | NA02 | NA13~NT0228~NT0217 |
|  | Tropical and Subtropical Coniferous Forests | NA03 | NA13~NA02~NT0228~NT0204~NT0310~NT0177~NT0176~NT0217 |
|  | Temperate Broadleaf and Mixed Forests | NA04 | NA08~NA07~NA06~NA05 |
|  | Temperate Coniferous Forests | NA05 | NA13~NA12~NA11~NA08~NA07~NA06~NA04~NT0904~NT0164 |
|  | Boreal Forests-Taiga | NA06 | NA11~NA08~NA05~NA04 |
|  | Tropical and subtropical grasslands, savannas, and Shrublands | NA07 | NA13~NA08~NA05~NA04 |
|  | Temperate Grasslands, Savannas, and Shrublands | NA08 | NA13~NA12~NA06~NA05~NA04 |
|  | Tundra | NA11 | NA06~NA05 |
|  | Mediterranean Forests, Woodlands, and Scrub | NA12 | NA13~NA08~NA05 |
|  | Deserts and Xeric Shrublands | NA13 | NA12~NA08~NA07~NA05~NA03~NA02~NT0176~NT0310~NT0204~NT0228 |
| **Neotropics** |  |  |  |
|  | Tropical and Subtropical Moist Broadleaf Forests | NT01 | NT02~NT03~NT07~NT09~NT10~NT13~NA0303~NA1311~NA1312 |
|  | Tropical and Subtropical Dry Broadleaf Forests | NT02 | NT01~NT03~NT07~NT09~NT10~NT13~NA1302~NA0302 |
|  | Tropical and Subtropical Coniferous Forests | NT03 | NT01~NT02~NA1302~NA0303 |
|  | Temperate Broadleaf and Mixed Forests | NT04 | NT10~NT12 |
|  | Tropical and subtropical grasslands, savannas, and Shrublands | NT07 | NT01~NT02~NT08~NT09~NT10~NT13 |
|  | Temperate Grasslands, Savannas, and Shrublands | NT08 | NT04~NT07~NT09~NT10 |
|  | Flooded Grasslands and Savannas | NT09 | NT01~NT02~NT07~NT08~NA0529 |
|  | Montane Grasslands and Shrublands | NT10 | NT01~NT02~NT04~NT07~NT08~NT12~NT13 |
|  | Mediterranean Forests, Woodlands, and Scrub | NT12 | NT04~NT09~NT13 |
|  | Deserts and Xeric Shrublands | NT13 | NT01~NT02~NT07~NT10~NT12 |
| **Oceania** |  |  |  |
|  | Tropical and Subtropical Moist Broadleaf Forests | OC01 | OC02~OC07 |
|  | Tropical and Subtropical Dry Broadleaf Forests | OC02 | OC01~OC07 |
| Tropical and subtropical grasslands, savannas, and shrublands | OC07 | OC02~OC01 |
| **Palearctic** |  |  |
| Tropical and Subtropical Moist Broadleaf Forests | PA01 | IM0118~IM0149~IM0137~PA04~PA05 |
| Temperate Broadleaf and Mixed Forests | PA04 | PA05~PA06~PA08~PA09~PA10~PA12~IM0118 |
| Temperate Coniferous Forests | PA05 | PA04~PA06~PA08~PA10~PA12~IM0140~IM0137~IM0402 |
| Boreal Forests-Taiga | PA06 | PA04~PA05~PA11 |
| Temperate Grasslands, Savannas, and Shrublands | PA08 | PA04~PA05~PA10~PA13 |
| Flooded Grasslands and Savannas | PA09 | PA04~PA08 |
| Montane Grasslands and Shrublands | PA10 | PA05~PA08~PA13 |
| Tundra | PA11 | PA06 |
| Mediterranean Forests, Woodlands, and Scrub | PA12 | PA04~PA05~PA13 |
| Deserts and Xeric Shrublands | PA13 | AT1303~AT1320~AT1306~AT1321~AT1302~AT0801~AT0713~PA08~PA10~PA12~IM1303~IM1302 |

**Figure 3:**
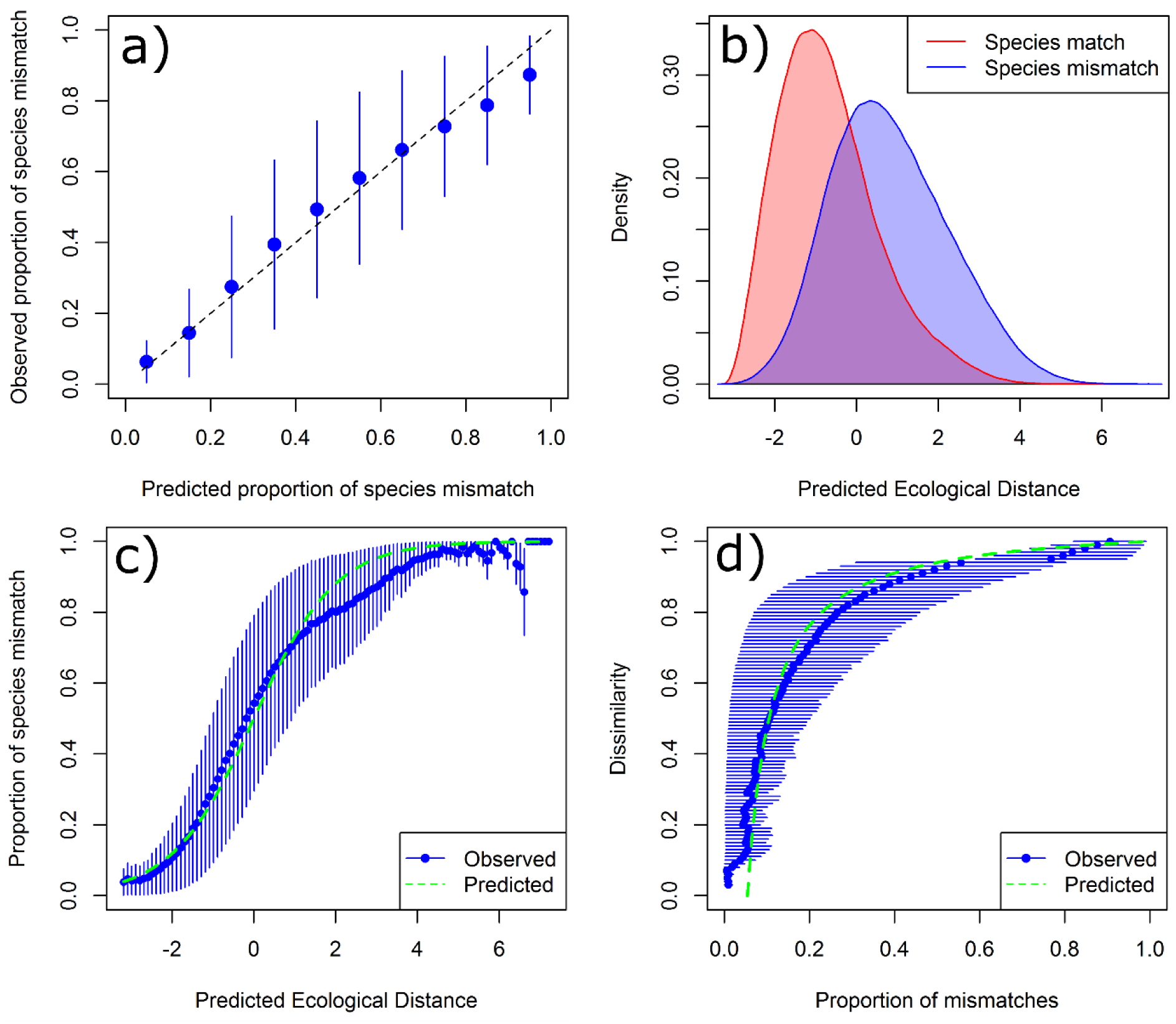
Example model fitting and validation outputs showing: (a) the observed proportion of species mismatches for 10 bins along the predicted proportion of species mismatches from the model. (b) the density of observed species matches and mismatches along the predicted ecological distance (in logit space) from the model. (c) the observed (blue) and predicted (green) proportion of species mismatches for 100 bins along the predicted ecological distance (in logit space). (d) comparison of observed (blue) and predicted (green) compositional dissimilarity against the proportion of mismatches. All error bars show the observed proportion of matches and mismatches ± the variance.

This allowed consideration of major biogeographic discontinuities between realms and potential variation in the response of different species assemblages between biomes. However biomes were, as far as possible, not treated as closed systems. Model fitting drew on biological and environmental data from both the core biome and adjacent biomes within the same realm, and from adjacent ecoregions in neighbouring realms where the realm boundary was considered porous to the movement of species (e.g. the Nearctic/Neotropics divide in Florida) (Table 3). Models were then fitted to a combination of pairs of species observations, in which 50% of the pairs consisted of observations exclusively drawn from within the target bio-realm and the other 50% paired an observation from within the bio-realm with an observation from one of the buffering regions (Table 3).

We sampled a maximum of 1.5 million unique pairs of species observations to fit the model for each combination of broad biological group (vascular plants, invertebrates, vertebrates) and bio-realm. Each pair of species observations was always drawn from within the same taxonomic sub-group, rather than between sub-groups (See Table 1 for groupings). The 1.5 million pairs of observations were drawn as evenly as possible from the sub-groups within each broad biological groups – e.g. for vertebrates, an equal number of observation pairs were drawn from the data for mammals, birds, reptiles and amphibians. Additionally, to maintain a reasonable level of data independence, and to avoid a small number of species observations unduly influencing the fitting of any model, each individual species observation was only used a maximum of 10 times in the sampling of observation pairs.

Our models were fitted using three I-spline basis functions for each environmental covariate (with the exception of bare ground which used a single linear function due to it essentially being a binary variable) with knots placed at the 0, 0.5 and 1 quantiles of the distribution of values for that covariate within each target bio-realm. Where the environmental envelope of samples taken from buffering regions lay outside the envelope for the target biome, additional knots were added at the outer limits, resulting in 3, 4 or 5 knots depending on the structure of the data. This allowed the description of spatial turnover in species composition to be tuned primarily to the target bio-realm while also accounting for extensions of these patterns into neighbouring regions. We regarded the latter as being of particular importance for any subsequent use of these models to project impacts of climate change on biodiversity distribution.

As indicated above, the effect of major biogeographic discontinuities on spatial turnover in species composition was accounted for by fitting separate models for different bio-realms. The effect of geographic separation on compositional turnover at finer spatial scales was then addressed by including straight-line geographic distance between species observations as an additional covariate in models fitted within each bio-realm. Observed correlation between geographic distance and compositional turnover can result not only from the effects of geographic isolation *per se*, but also from high levels of spatial autocorrelation exhibited by many environmental variables (Warren *et al.*, 2014). We therefore took considerable care to avoid geographic distance overwhelming, or masking, the effects of more direct environmental predictors within our models. We achieved this by fitting each model in a two-staged procedure; firstly fitting to the *a priori* selected set of environmental predictor variables alone, and then refitting the model with the combined effect of the environmental covariates now fixed as an offset, thereby allowing the addition of geographic distance to describe only variation not already accounted for by the environmental covariates. The effect of geographic distance was fitted using a linear function, rather than the complex splined functions used for the environmental covariates.

To aid in model fitting efficiency, the number of matching species observations used was up-weighted to provide a sample set that included a 1:1 ratio of species matches and mismatches. However, this up-weighting has the effect of disrupting the relationship between our modelled property (*p_i,j_*) and compositional turnover. Following model fitting the predicted probabilities of obtaining a species mismatch (*p*_*i,j*_) were corrected using:

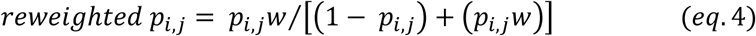

where (*w*_*m*_) represents a pre-calculated weighting value derived from the ratio of species mismatches and species matches from a completely random sample of species observation pairs for each species x bio-realm combination (𝑚). Adding *eq.* 4 into *eq.* 3 we then use:

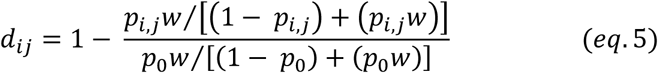

to derive our estimate of compositional turnover.

### Example model outputs

Our fitted models greatly refine the mapping of beta-diversity patterns relative to more traditional surrogates such as ecoregions (Fig. 4a). By working at a biologically-relevant spatial resolution, and allowing biological composition to vary continuously across environmental and geographic space at this resolution, we are able to predict finer-scaled patterns of variation within ecoregions, or other large discrete units, and across the boundaries of these units (Fig. 4b). In a recent evaluation of the performance of GDM in mapping beta-diversity patterns across the Australian continent, Ware *et al.* (2018) demonstrated that GDM models fitted to scattered presence-only data on species occurrences, of the type employed here in our global modelling, achieved significantly better concordance with actual patterns of spatial variation in biological composition, based on independent test datasets for plants, invertebrates and vertebrates, than that achieved by best-available mapping of bioregions and major vegetation types.

**Figure 4:**
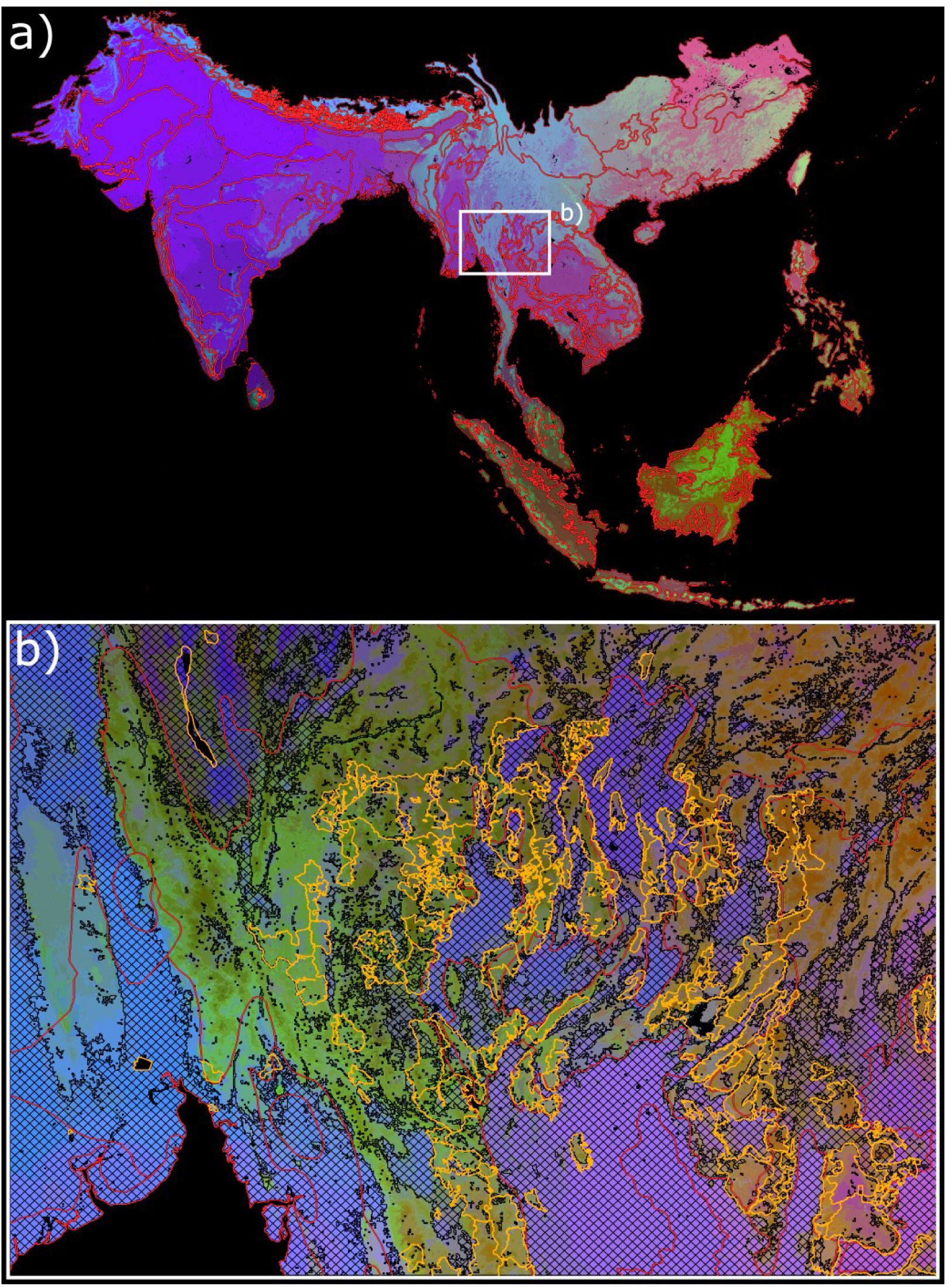
Visualisation of the spatial patterns in beta-diversity being estimated using the BILBI modelling framework for vertebrates in the Indo-Malaysian realm (a). The colours across the surface represent the level of similarity estimated between regions with similarity depicted via similar colours. Red lines represent the WWF ecoregional boundaries described by Olson *et al.* (2001). Zoomed map (b) shows modelled results for vertebrate communities with cleared areas covered with black hatching and protected areas shown in the green polygons. Note: RGB colour ramps have been stretched across each of the two different analysis domains to maximise the visual discrimination shown in each map.

When mapping of habitat loss and protected-area boundaries are superimposed over our modelled patterns of compositional variation, simple visual inspection suggests strong levels of covariance between these layers. In other words habitat loss and protection are not distributed randomly in relation to finer-scaled compositional variation, but are instead biased towards particular subsets of this variation. This level of covariance opens up considerable potential to derive protected-area and habitat indicators globally which account more effectively for biases in the distribution of habitat protection and loss playing out at a resolution below that of ecoregions (Fig 4b). We outline current initiatives tapping into this potential in the following section.

While we prefer to present results of our modelling as continuous patterns of compositional turnover, and to employ these continuous predictions directly in any subsequent assessment, we also recognise that some applications may be constrained to working with biodiversity surrogates taking the form of a discrete classification. For such applications there is potential to derive discrete classes (i.e. ‘ecosystems’ or ‘biologically-scaled environmental domains’) from our modelling of continuous patterns of compositional turnover, through numerical classification (Fig. 5a). This involves using the predicted compositional similarity between pairs of grid-cells to numerically cluster similar cells into discrete classes (for further explanation of this approach seeFerrier *et al.*, 2007; Leathwick *et al.*, 2011). Classifications such as these can be generated for a hierarchy of different spatial domains ranging from, for example, whole realms or bio-realms (Fig. 5a) through to further subdivision of any given class within this larger domain (Fig. 5b). Relative to approaches that rely on classifying patterns based on environmental variables alone (e.g. Sayre *et al.*, 2014), classifications derived from this approach have the benefit of incorporating best-available biological information through the scaling of environmental space based on modelled patterns in compositional turnover. In an evaluation of the performance of a GDM-based classification of New Zealand’s river and stream system, Leathwick *et al.* (2011) found that this approach achieved significantly better discrimination of independently observed biological patterns than did either rule-based or numerical classifications based on environmental variables alone.

**Figure 5:**
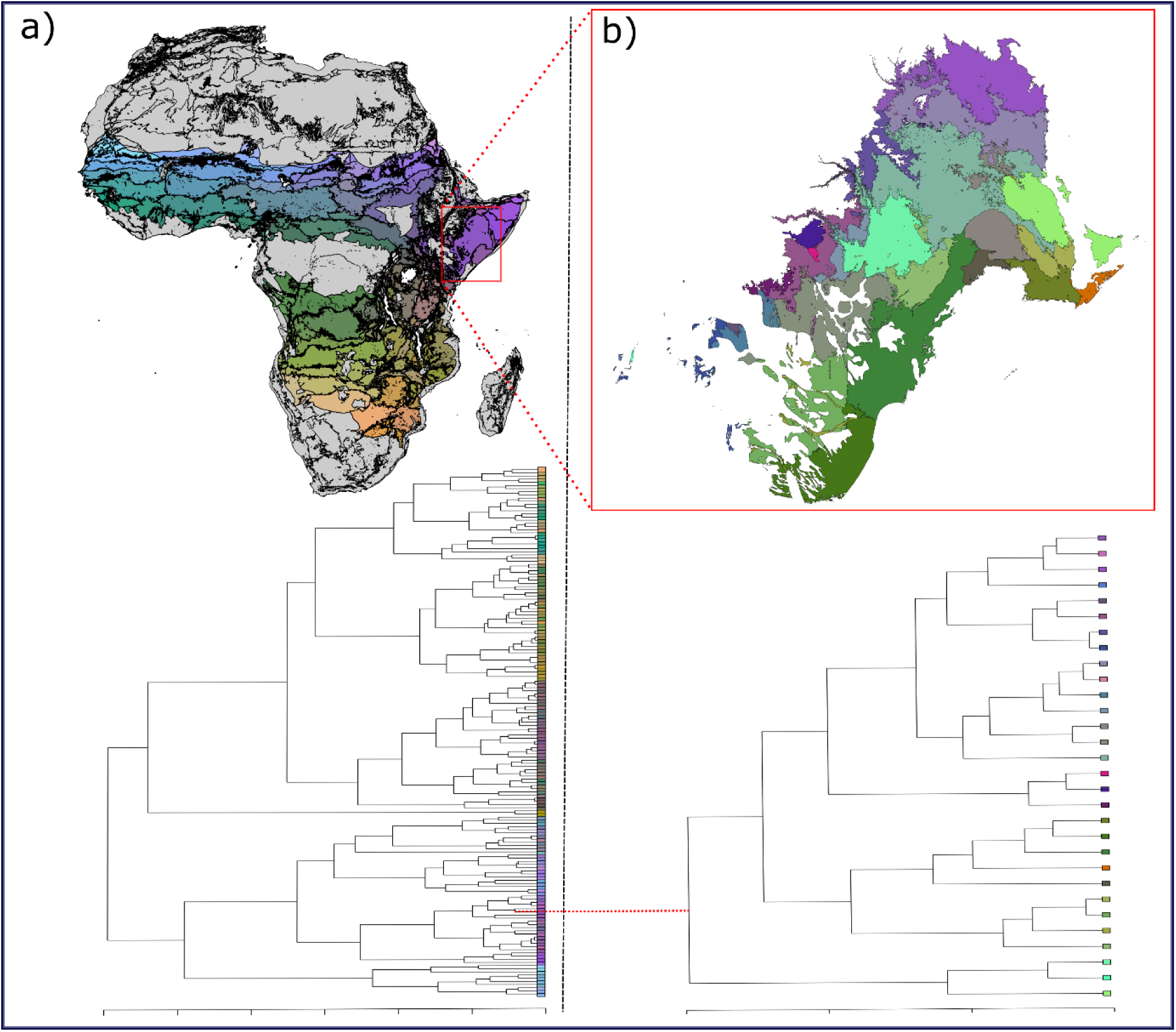
Numerical classification of the BILBI derived continuous predictions of turnover in vertebrate communities for the entire African continent (a) with a classification for the African tropical savannah biorealm nested inside (shown in colour, with each class coloured by its similarity to other classes). Additional classification of a single unit from the African tropical savannah biorealm is shown in (b). The relationships between each class are shown in the dendrograms below each map. Coloured leaves on the dendrograms relate to the colours of each class on the maps above.

### EMERGING APPLICATIONS TO GLOBAL BIODIVERSITY ASSESSMENT

As depicted in Fig. 2 our modelling of spatial patterns in the distribution of biodiversity within BILBI is intended to inform global biodiversity assessment through two subsequent pathways of analysis: 1) generation of indicators of past-to-present change in biodiversity based on observed changes in habitat condition and protected-area coverage; and 2) projection of potential future change in biodiversity as a consequence of alternative scenarios of global change (particularly changes in land-use and climate) and associated policy options. Considerable progress has already been made in applying the BILBI modelling framework to both of these activities over the past few years. While we here provide a brief overview of this work, detailed description of the techniques employed in, and the results obtained from, these applications are beyond the scope of this paper, and will be covered in a number of forthcoming publications.

The BILBI framework has been used to generate two global indicators for reporting progress against CBD Aichi Targets. Both of these are now included in the CBD’s official list of recognised indicators (https://www.cbd.int/doc/decisions/cop-13/cop-13-dec-28-en.pdf), and in the suite of indicators coordinated and promoted by the Biodiversity Indicators Partnership (https://www.bipindicators.net/). The first of these two indicators – the Protected Area Representativeness Index – assesses the extent to which terrestrial protected areas are “ecologically representative”, in accordance with Aichi Target 11 (https://www.bipindicators.net/indicators/protected-area-representativeness-index-parc-representativeness). This assessment is performed at a much finer ecological and spatial resolution than that typically employed in other assessments of protected-area representativeness. This is achieved by combining BILBI’s modelling of spatial turnover in biodiversity composition with the World Database on Protected Areas (UNEP & WCMC, 2016), and the analytical approach to assessing representativeness using continuous predictions of compositional turnover described by Ferrier *et al.* (2004) and Allnutt *et al.* (2008)

The second indicator – the Biodiversity Habitat index – is intended to add value to existing assessments of the “rate of loss … of all natural habitats, including forests”, under Aichi Target 5, by translating the observed spatial distribution of habitat loss and degradation into expected impacts on retention of terrestrial biodiversity (https://www.bipindicators.net/indicators/biodiversity-habitat-index). This indicator is generated using the same analytical approach underpinning the Protected Area Representativeness Index, but now combining BILBI’s compositional-turnover modelling with best-available data on habitat condition in place of protected-area coverage. Initially the indicator was derived for forest biomes only, using spatially-explicit data on forest loss from the Global Forest Change dataset (Hansen *et al.*, 2013). However it is now being expanded to cover all terrestrial biomes across the planet by estimating condition through an extension of the Hoskins *et al.* (2016) statistical downscaling of coarse-resolution land-use data using 30-arcsecond environmental and remotely-sensed land-cover covariates. This work has adapted the Hoskins *et al.* (2016) approach to employ Version 2, in place of Version 1, of the Land Use Harmonisation product, thereby generating downscaled estimates of 12, rather than the original five, land-use classes (http://luh.umd.edu/), and MODIS Vegetation Continuous Fields (http://glcf.umd.edu/data/vcf/) as remote-sensing covariates in place of discrete land-cover classes. Applying this downscaling approach across multiple years provides an effective means of translating observed changes in remote-sensing covariates into estimated changes in the proportions of land-use classes occurring in each and every 30-arcsecond terrestrial grid-cell on the planet. These proportions can then, in turn, being translated into an estimate of habitat condition, for any given cell in any given year, using coefficients derived from global meta-analyses of land-use impacts on local retention of species diversity undertaken, for example, by the PREDICTS project (Hudson *et al.*, 2014; Newbold *et al.*, 2016).

In addition to generating the above indicators of past-to-present change, the BILBI framework is also being used to project potential future change in biodiversity as a consequence of alternative scenarios of global change and associated policy options. BILBI has already been employed, alongside a range of other biodiversity and ecosystem modelling approaches, in two major multi-model scenario analyses – one assessing the potential biodiversity and ecosystem-service impacts of selected combinations of land-use trajectories (based on Shared Socio-economic Pathways) and climate trajectories (based on Representative Concentration Pathways) (Kim *et al.*, 2018); and the other assessing policy options to reverse ongoing biodiversity loss resulting from detrimental changes in land use (Leclere *et al.*, 2018). To project the potential effects of climate change on beta-diversity patterns space-for-time substitution is used to predict the level of compositional change expected over time, as a function of BILBI’s compositional-turnover models describing how species composition changes spatially along present-day environmental gradients (for further detail on GDM-based projection of compositional change under climate change see: Fitzpatrick *et al.*, 2011; Blois *et al.*, 2013). Fine-scaled projection of changes in habitat condition is achieved by linking statistical downscaling of present-day land use (described above) with coarse-resolution scenarios of land-use change. The species-area relationship (SAR) is then used to predict the proportion of species expected to persist over the longer term, as a function of the effective proportion of habitat remaining. In contrast to other SAR-based approaches, which work with discrete environmental classes or ecosystem types, this approach applies the SAR to biologically-scaled environments varying continuously across space and time (for further detail of this particular approach see: Ferrier *et al.*, 2004; Allnutt *et al.*, 2008).

Two other broad areas of potential application of the BILBI modelling framework are worth noting at this point. GDM-based modelling of compositional turnover has been applied previously in both of these contexts, but only at sub-global scales (Ferrier *et al.*, 2007). Establishment of the BILBI framework now opens up new opportunities to extend these applications globally. The first of these would involve using the scaled multidimensional environmental space, resulting from the fitting of GDM models within BILBI, to extrapolate geographical distributions of individual species as a function of the density of occurrence records for any given species across this environmental space. When (Elith *et al.*, 2006) tested this approach by applying GDM, coupled with a simple kernel-regression technique, to data from six study-regions around the world they achieved a level of predictive performance very similar to that of MaxEnt (which later became the world’s most widely applied SDM technique) and markedly better than that achieved by most of the other SDM techniques evaluated in that study. Considerable potential now exists to employ the GDM models already fitted within the BILBI framework as a foundation for extrapolating the potential global distribution of any species with a sufficient number of occurrence records from the > 400,000 species of plants, invertebrates and vertebrates contributing data to these GDMs.

The second of these potential applications would involve using the fitted GDM models within BILBI to help assess the adequacy of biological sampling across environmental and geographical space, and to identify gaps in this coverage to help prioritise future survey effort. Ferrier (2002) outlined an iterative strategy for survey gap-analysis and prioritisation which couples GDM modelling with p-median analysis, an operations-research technique originally adapted for use in conservation biology by Faith and Walker (1996). This strategy prioritises locations which can best fill gaps in the coverage of existing survey sites across a GDM-scaled multidimensional environmental and geographical space fitted to the biological data from those existing sites. Data collected at these new sites can then be used to test and refit the GDM, thereby enabling iterative refinement of both the modelling itself, and the prioritisation of remaining gaps in biological data coverage. While the potential of this survey gap-analysis strategy has already been demonstrated globally using an unscaled environmental-geographical space, as part of the trial GBIF-MAPA application (Flemons *et al.*, 2007), implementation of the fully coupled GDM/p-median approach has, until now, been limited to selected regions and taxa (e.g Ferrier *et al.*, 2007; Bell *et al.*, 2014). The establishment of global GDM models within the BILBI framework has therefore now opened up an unprecedented opportunity to apply this approach across the entire terrestrial surface of the planet. This would help to direct future biological survey and collection efforts to maximise not only the environmental and geographical coverage of occurrence records for known species, but also the likelihood of discovering new species previously unknown to science, especially within lesser-studied, yet hyper-diverse, taxa.

## CONCLUSION

The BILBI modelling framework offers a means of making more effective use of scattered occurrence data for highly-diverse biological groups to map fine-scaled patterns in the distribution of biodiversity worldwide – including robust extrapolation of expected patterns across poorly-sampled regions, even where the particular species occurring in these regions are unknown or unsurveyed. This capability is intended to complement, rather than compete with, other approaches to global biodiversity assessment, including those employing discrete land classifications (e.g. ecoregions, ecosystem types) and those focussed on individual species within better-known biological groups, or on species of particular conservation concern.

While the initial implementation of BILBI relies strongly on modelling of beta diversity patterns using one particular technique (GDM) we have purposely designed the overall framework to be sufficiently generic and flexible to allow incremental refinement and addition of modelling techniques and datasets into the future. As noted earlier, work is already proceeding to incorporate mapping of alpha-diversity patterns, alongside the existing mapping of beta diversity, by extending our modelling approach to simultaneously model both species richness and compositional turnover from presence-only data. Also high on our list of priorities for model enhancement is to more rigorously account for the effects of biogeographical barriers on compositional turnover, at multiple scales. In its current form BILBI accounts for such effects only by assuming complete distinctiveness in species composition between major biogeographical realms, and through inclusion of straight-line spatial separation (or ‘isolation by distance’) as an explanatory variable, alongside environmental predictors, in the fitting of GDMs. Considerable potential exists to refine this approach by incorporating measures of ecological isolation (e.g. isolation of islands by intervening ocean, or isolation of mountains by intervening lowlands) into the modelling (Ferrier *et al.*, 2007).

There is also considerable potential to refine our modelling of biodiversity patterns within BILBI through incorporation of new and emerging sources of environmental and biological data. This includes advances in the use of remote sensing to derive global environmental surfaces tailored specifically to the needs of biodiversity modelling (e.g. Wilson & Jetz, 2016). In terms of biological data, one of our highest priorities is to augment our existing focus on species-occurrence data (i.e. presence-only records) with greater use of inventory datasets, particularly regional and global compilations of survey-plot data for plants (e.g. Franklin *et al.*, 2017). While the geographical coverage of such datasets is often patchy relative to species-occurrence datasets, they offer valuable potential to independently test the performance of existing BILBI models in predicting compositional turnover between rigorously-sampled sites, and to compare this performance with that of other modelling techniques (e.g. SDMs) and biodiversity surrogates (e.g. mapped ecoregions or ecosystems). Another priority is to make better use of information on phylogenetic relationships between species, where available for particular biological groups, to extend our current modelling of beta diversity to account for turnover in phylogenetic composition, rather than simply taxonomic composition (Rosauer *et al.*, 2014).

## Supporting information

R code detailing analytical approach

## References

Allnutt, T.F., Ferrier, S., Manion, G., Powell, G.V.N., Ricketts, T.H., Fisher, B.L., Harper, G.J., Irwin, M.E., Kremen, C., Labat, J.N., Lees, D.C., Pearce, T.A. & Rakotondrainibe, F. (2008) A method for quantifying biodiversity loss and its application to a 50-year record of deforestation across Madagascar. Conservation Letters, 1, 173–181.

Amatulli, G., Domisch, S., Tuanmu, M.-N., Parmentier, B., Ranipeta, A., Malczyk, J. & Jetz, W. (2018) A suite of global, cross-scale topographic variables for environmental and biodiversity modeling. Scientific Data, 5, 180040.

Beck, J., Holloway, J.D., Schwanghart, W. & Orme, D. (2013) Undersampling and the measurement of beta diversity. Methods in Ecology and Evolution, 4, 370–382.

Bell, K.L., Heard, T.A., Manion, G., Ferrier, S. & van Klinken, R.D. (2014) Characterising the phytophagous arthropod fauna of a single host plant species: assessing survey completeness at continental and local scales. Biodiversity and Conservation, 23, 2985–3003.

Blois, J.L., Williams, J.W., Fitzpatrick, M.C., Ferrier, S., Veloz, S.D., He, F., Liu, Z.Y., Manion, G. & Otto-Bliesner, B. (2013) Modeling the climatic drivers of spatial patterns in vegetation composition since the Last Glacial Maximum. Ecography, 36, 460–473.

Butchart, S.H.M., Clarke, M., Smith, R.J., Sykes, R.E., Scharlemann, J.P.W., Harfoot, M., Buchanan, G.M., Angulo, A., Balmford, A., Bertzky, B., Brooks, T.M., Carpenter, K.E., Comeros-Raynal, M.T., Cornell, J., Ficetola, G.F., Fishpool, L.D.C., Fuller, R.A., Geldmann, J., Harwell, H., Hilton‐Taylor, C., Hoffmann, M., Joolia, A., Joppa, L., Kingston, N., May, I., Milam, A., Polidoro, B., Ralph, G., Richman, N., Rondinini, C., Segan, D.B., Skolnik, B., Spalding, M.D., Stuart, S.N., Symes, A., Taylor, J., Visconti, P., Watson, J.E.M., Wood, L. & Burgess, N.D. (2015) Shortfalls and Solutions for Meeting National and Global Conservation Area Targets. Conservation Letters, 8, 329–337.

Calderón-Patrón, J.M., Goyenechea, I., Ortiz-Pulido, R., Castillo-Cerón, J., Manriquez, N., Ramírez-Bautista, A., Rojas-Martínez, A.E., Sánchez-Rojas, G., Zuria, I. & Moreno, C.E. (2016) Beta Diversity in a Highly Heterogeneous Area: Disentangling Species and Taxonomic Dissimilarity for Terrestrial Vertebrates. PLOS ONE, 11, e0160438.

CBD (2010) Strategic plan for biodiversity. In:

Citroen, S., Kempinski, J. & Cullen, Z. (2016) Life after COP21: what does the Paris Agreement mean for forests and biodiversity conservation?. Oryx, 50, 201–202.

D’Amen, M., Rahbek, C., Zimmermann, N.E. & Guisan, A. (2017) Spatial predictions at the community level: from current approaches to future frameworks. Biological Reviews, 92, 169–187.

Danielson, J.J.G. & Dean, B. (2011) Global Multi-resolution Terrain Elevation Data 2010 (GMTED2010): U.S. Geological Survey Open-File Report 2011-1073. In: (ed. U.S.D.O.T. Interior), p. 26. U.S. Geological Survey, Reston, Virginia.

Danielson, J.J.G., Dean B. (2011) Global Multi-resolution Terrain Elevation Data 2010 (GMTED2010): U.S. Geological Survey Open-File Report 2011-1073. In: (ed. U.S.D.O.T. Interior), p. 26. U.S. Geological Survey

Elith, J. & Leathwick, J.R. (2009) Species Distribution Models: Ecological Explanation and Prediction Across Space and Time. Annual Review of Ecology, Evolution, and Systematics, 40, 677–697.

Elith, J., Graham, C.H., Anderson, R.P., Dudik, M., Ferrier, S., Guisan, A., Hijmans, R.J., Huettmann, F., Leathwick, J.R., Lehmann, A., Li, J., Lohmann, L.G., Loiselle, B.A., Manion, G., Moritz, C., Nakamura, M., Nakazawa, Y., Overton, J.M., Peterson, A.T., Phillips, S.J., Richardson, K., Scachetti-Pereira, R., Schapire, R.E., Soberon, J., Williams, S., Wisz, M.S. & Zimmermann, N.E. (2006) Novel methods improve prediction of species’ distributions from occurrence data. Ecography, 29, 129–151.

Faith, D.P. & Walker, P.A. (1996) Environmental diversity: on the best-possible use of surrogate data for assessing the relative biodiversity of sets of areas. Biodiversity & Conservation, 5, 399–415.

FAO/IIASA/ISRIC/ISS-CAS/JRC (2012) Harmonised World Soil Database v 1.2. In: (ed. R. Fao, Italy & Iiasa, Laxenburg, Austria)

Ferrier, S. (2002) Mapping spatial pattern in biodiversity for regional conservation planning: Where to from here?. Systematic Biology, 51, 331–363.

Ferrier, S. (2011) Extracting More Value from Biodiversity Change Observations through Integrated Modeling. Bioscience, 61, 96–97.

Ferrier, S. & Guisan, A. (2006) Spatial modelling of biodiversity at the community level. Journal of Applied Ecology, 43, 393–404.

Ferrier, S., Manion, G., Elith, J. & Richardson, K. (2007) Using generalized dissimilarity modelling to analyse and predict patterns of beta diversity in regional biodiversity assessment. Diversity and Distributions, 13, 252–264.

Ferrier, S., Powell, G.V.N., Richardson, K.S., Manion, G., Overton, J.M., Allnutt, T.F., Cameron, S.E., Mantle, K., Burgess, N.D., Faith, D.P., Lamoreux, J.F., Kier, G., Hijmans, R.J., Funk, V.A., Cassis, G.A., Fisher, B.L., Flemons, P., Lees, D., Lovett, J.C. & Van Rompaey, R.S.A.R. (2004) Mapping more of terrestrial biodiversity for global conservation assessment. Bioscience, 54, 1101–1109.

Fitzpatrick, M.C., Sanders, N.J., Ferrier, S., Longino, J.T., Weiser, M.D. & Dunn, R. (2011) Forecasting the future of biodiversity: a test of single- and multi-species models for ants in North America. Ecography, 34, 836–847.

Flemons, P., Guralnick, R., Krieger, J., Ranipeta, A. & Neufeld, D. (2007) A web-based GIS tool for exploring the world’s biodiversity: The Global Biodiversity Information Facility Mapping and Analysis Portal Application (GBIF-MAPA). Ecological Informatics, 2, 49–60.

Francis, A.P. & Currie, D.J. (1998) Globl patterns of tree species richness in moise forests: another look. Oikos, 81, 598–602.

Franklin, J., Serra-Diaz, J.M., Syphard, A.D. & Regan, H.M. (2017) Big data for forecasting the impacts of global change on plant communities. Global Ecology and Biogeography, 26, 6–17.

GBIF.org (20 March 2014) Custom GBIF data exprt. In: https://doi.org/10.15468/cdl.qhsa6c

Hansen, M.C., Potapov, P.V., Moore, R., Hancher, M., Turubanova, S.A., Tyukavina, A., Thau, D., Stehman, S.V., Goetz, S.J., Loveland, T.R., Kommareddy, A., Egorov, A., Chini, L., Justice, C.O. & Townshend, J.R.G. (2013) High-Resolution Global Maps of 21st-Century Forest Cover Change. Science, 342, 850–853.

Hengl, T., de Jesus, J.M., MacMillan, R.A., Batjes, N.H., Heuvelink, G.B.M., Ribeiro, E., Samuel-Rosa, A., Kempen, B., Leenaars, J.G.B., Walsh, M.G. & Gonzalez, M.R. (2014) SoilGrids1km — Global Soil Information Based on Automated Mapping. PLOS ONE, 9, e105992.

Hengl, T.K., M (2013) TWISRE3: SAGA GIS Topographic wetness index. In: (ed. Worldgrids)

Hijmans, R.J., Cameron, S.E., Parra, J.L., Jones, P.G. & Jarvis, A. (2005) Very high resolution interpolated climate surfaces for global land areas. International Journal of Climatology, 25, 1965–1978.

Hoskins, A.J., Bush, A., Gilmore, J., Harwood, T., Hudson, L.N., Ware, C., Williams, K.J. & Ferrier, S. (2016) Downscaling land-use data to provide global 30″ estimates of five land-use classes. Ecology and Evolution, 6, 3040–3055.

Hudson, L.N., Newbold, T., Contu, S., Hill, S.L.L., Lysenko, I., De Palma, A., Phillips, H.R.P., Senior, R.A., Bennett, D.J., Booth, H., Choimes, A., Correia, D.L.P., Day, J., Echeverría-Londoño, S., Garon, M., Harrison, M.L.K., Ingram, D.J., Jung, M., Kemp, V., Kirkpatrick, L., Martin, C.D., Pan, Y., White, H.J., Aben, J., Abrahamczyk, S., Adum, G.B., Aguilar-Barquero, V., Aizen, M.A., Ancrenaz, M., Arbeláez-Cortés, E., Armbrecht, I., Azhar, B., Azpiroz, A.B., Baeten, L., Báldi, A., Banks, J.E., Barlow, J., Batáry, P., Bates, A.J., Bayne, E.M., Beja, P., Berg, Å., Berry, N.J., Bicknell, J.E., Bihn, J.H., Böhning-Gaese, K., Boekhout, T., Boutin, C., Bouyer, J., Brearley, F.Q., Brito, I., Brunet, J., Buczkowski, G., Buscardo, E., Cabra-García, J., Calviño-Cancela, M., Cameron, S.A., Cancello, E.M., Carrijo, T.F., Carvalho, A.L., Castro, H., Castro-Luna, A.A., Cerda, R., Cerezo, A., Chauvat, M., Clarke, F.M., Cleary, D.F.R., Connop, S.P., D’Aniello, B., da Silva, P.G., Darvill, B., Dauber, J., Dejean, A., Diekötter, T., Dominguez-Haydar, Y., Dormann, C.F., Dumont, B., Dures, S.G., Dynesius, M., Edenius, L., Elek, Z., Entling, M.H., Farwig, N., Fayle, T.M., Felicioli, A., Felton, A.M., Ficetola, G.F., Filgueiras, B.K.C., Fonte, S.J., Fraser, L.H., Fukuda, D., Furlani, D., Ganzhorn, J.U., Garden, J.G., Gheler-Costa, C., Giordani, P., Giordano, S., Gottschalk, M.S., Goulson, D., Gove, A.D., Grogan, J., Hanley, M.E., Hanson, T., Hashim, N.R., Hawes, J.E., Hébert, C., Helden, A.J., Henden, J.-A., Hernández, L., Herzog, F., Higuera-Diaz, D., Hilje, B., Horgan, F.G., Horváth, R., Hylander, K., Isaacs-Cubides, P., Ishitani, M., Jacobs, C.T., Jaramillo, V.J., Jauker, B., Jonsell, M., Jung, T.S., Kapoor, V., Kati, V., Katovai, E., Kessler, M., Knop, E., Kolb, A., Kőrösi, Á., Lachat, T., Lantschner, V., Le Féon, V., LeBuhn, G., Légaré, J.-P., Letcher, S.G., Littlewood, N.A., López-Quintero, C.A., Louhaichi, M., Lövei, G.L., Lucas-Borja, M.E., Luja, V.H., Maeto, K., Magura, T., Mallari, N.A., Marin-Spiotta, E., Marshall, E.J.P., Martínez, E., Mayfield, M.M., Mikusinski, G., Milder, J.C., Miller, J.R., Morales, C.L., Muchane, M.N., Muchane, M., Naidoo, R., Nakamura, A., Naoe, S., Nates-Parra, G., Navarrete Gutierrez, D.A., Neuschulz, E.L., Noreika, N., Norfolk, O., Noriega, J.A., Nöske, N.M., O’Dea, N., Oduro, W., Ofori-Boateng, C., Oke, C.O., Osgathorpe, L.M., Paritsis, J., Parra-H, A., Pelegrin, N., Peres, C.A., Persson, A.S., Petanidou, T., Phalan, B., Philips, T.K., Poveda, K., Power, E.F., Presley, S.J., Proença, V., Quaranta, M., Quintero, C., Redpath-Downing, N.A., Reid, J.L., Reis, Y.T., Ribeiro, D.B., Richardson, B.A., Richardson, M.J., Robles, C.A., Römbke, J., Romero-Duque, L.P., Rosselli, L., Rossiter, S.J., Roulston, T.a.H., Rousseau, L., Sadler, J.P., Sáfián, S., Saldaña-Vázquez, R.A., Samnegård, U., Schüepp, C., Schweiger, O., Sedlock, J.L., Shahabuddin, G., Sheil, D., Silva, F.A.B., Slade, E.M., Smith-Pardo, A.H., Sodhi, N.S., Somarriba, E.J., Sosa, R.A., Stout, J.C., Struebig, M.J., Sung, Y.-H., Threlfall, C.G., Tonietto, R., Tóthmérész, B., Tscharntke, T., Turner, E.C., Tylianakis, J.M., Vanbergen, A.J., Vassilev, K., Verboven, H.A.F., Vergara, C.H., Vergara, P.M., Verhulst, J., Walker, T.R., Wang, Y., Watling, J.I., Wells, K., Williams, C.D., Willig, M.R., Woinarski, J.C.Z., Wolf, J.H.D., Woodcock, B.A., Yu, D.W., Zaitsev, A.S., Collen, B., Ewers, R.M., Mace, G.M., Purves, D.W., Scharlemann, J.P.W. & Purvis, A. (2014) The PREDICTS database: a global database of how local terrestrial biodiversity responds to human impacts. Ecology and Evolution, 4, 4701–4735.

Hurlbert, A.H. & Jetz, W. (2007) Species richness, hotspots, and the scale dependence of range maps in ecology and conservation. Proceedings of the National Academy of Sciences, 104, 13384–13389.

Isaac, N.J.B. & Pocock, M.J.O. (2015) Bias and information in biological records. Biological Journal of the Linnean Society, 115, 522–531.

Jenkins, C.N., Pimm, S.L. & Joppa, L.N. (2013) Global patterns of terrestrial vertebrate diversity and conservation. Proceedings of the National Academy of Sciences, 110, E2602–E2610.

Jetz, W., McPherson, J.M. & Guralnick, R.P. (2012) Integrating biodiversity distribution knowledge: toward a global map of life. Trends in Ecology & Evolution, 27, 151–159.

Jones, M.M., Ferrier, S., Condit, R., Manion, G., Aguilar, S. & Perez, R. (2013) Strong congruence in tree and fern community turnover in response to soils and climate in central Panama. Journal of Ecology, 101, 506–516.

Kim, H., Rosa, I.M.D., Alkemade, R., Leadley, P., Hurtt, G., Popp, A., van Vuuren, D.P., Anthoni, P., Arneth, A., Baisero, D., Caton, E., Chaplin-Kramer, R., Chini, L., De Palma, A., Di Fulvio, F., Di Marco, M., Espinoza, F., Ferrier, S., Fujimori, S., Gonzalez, R.E., Gueguen, M., Guerra, C., Harfoot, M., Harwood, T.D., Hasegawa, T., Haverd, V., Havlík, P., Hellweg, S., Hill, S.L.L., Hirata, A., Hoskins, A.J., Janse, J.H., Jetz, W., Johnson, J.A., Krause, A., Leclère, D., Martins, I.S., Matsui, T., Merow, C., Obersteiner, M., Ohashi, H., Poulter, B., Purvis, A., Quesada, B., Rondinini, C., Schipper, A.M., Sharp, R., Takahashi, K., Thuiller, W., Titeux, N., Visconti, P., Ware, C., Wolf, F. & Pereira, H.M. (2018) A protocol for an intercomparison of biodiversity and ecosystem services models using harmonized land-use and climate scenarios. Geosci. Model Dev., 11, 4537–4562.

König, C., Weigelt, P., Kreft, H. & Baselga, A. (2017) Dissecting global turnover in vascular plants. Global Ecology and Biogeography, 26, 228–242.

Leathwick, J.R., Snelder, T., Chadderton, W.L., Elith, J., Julian, K. & Ferrier, S. (2011) Use of generalised dissimilarity modelling to improve the biological discrimination of river and stream classifications. Freshwater Biology, 56, 21–38.

Leclere, D., Obersteiner, M., Alkemade, R., Almond, R., Barrett, M., Bunting, G., Burgess, N., Butchart, S., Chaudhary, A., Cornell, S., De Palma, A., DeClerck, F., Di Fulvio, F., Di Marco, M., Doelman, J., Dürauer, M., Ferrier, S., Freeman, R., Fritz, S., Fujimori, S., Grooten, M., Harfoot, M., Harwood, T., Hasegawa, T., Havlik, P., Hellweg, S., Herrero, M., Hilbers, J., Hill, S., Hoskins, A., Humpenöder, F., Kram, T., Krisztin, T., Lotze-Campen, H., Mace, G., Matsui, T., Meyer, C., Nel, D., Newbold, T., Ohashi, H., Popp, A., Purvis, A., Schipper, A., Schmidt-Traub, G., Stehfest, E., Strassburg, B., Tabeau, A., Valin, H., van Meijl, H., van Vuuren, D., van Zeist, W., Visconti, P., Ware, C., Watson, J., Wu, W. & Young, L. (2018) Towards pathways bending the curve terrestrial biodiversity trends within the 21st century. In. IIASA

Lehner, B. & Doll, P. (2004) Development and validation of a global database of lakes, reservoirs and wetlands. Journal of Hydrology, 296, 1–22.

Londoño-Murcia, M.C., Tellez-Valdés, O. & Sánchez-Cordero, V. (2010) Environmental heterogeneity of World Wildlife Fund for Nature ecoregions and implications for conservation in Neotropical biodiversity hotspots. Environmental Conservation, 37, 116–127.

Merow, C., Wilson, A.M., Jetz, W. & Peres-Neto, P. (2017) Integrating occurrence data and expert maps for improved species range predictions. Global Ecology and Biogeography, 26, 243–258.

Meyer, C., Kreft, H., Guralnick, R. & Jetz, W. (2015) Global priorities for an effective information basis of biodiversity distributions. Nature Communications, 6, 8221.

Newbold, T., Hudson, L.N., Arnell, A.P., Contu, S., De Palma, A., Ferrier, S., Hill, S.L.L., Hoskins, A.J., Lysenko, I., Phillips, H.R.P., Burton, V.J., Chng, C.W.T., Emerson, S., Gao, D., Pask-Hale, G., Hutton, J., Jung, M., Sanchez-Ortiz, K., Simmons, B.I., Whitmee, S., Zhang, H., Scharlemann, J.P.W. & Purvis, A. (2016) Has land use pushed terrestrial biodiversity beyond the planetary boundary? A global assessment. Science, 353, 288–291.

Olson, D.M., Dinerstein, E., Wikramanayake, E.D., Burgess, N.D., Powell, G.V.N., Underwood, E.C., D’amico, J.A., Itoua, I., Strand, H.E., Morrison, J.C., Loucks, C.J., Allnutt, T.F., Ricketts, T.H., Kura, Y., Lamoreux, J.F., Wettengel, W.W., Hedao, P. & Kassem, K.R. (2001) Terrestrial Ecoregions of the World: A New Map of Life on Earth: A new global map of terrestrial ecoregions provides an innovative tool for conserving biodiversity. BioScience, 51, 933–938.

Pereira, H.M., Leadley, P.W., Proença, V., Alkemade, R., Scharlemann, J.P.W., Fernandez-Manjarrés, J.F., Araújo, M.B., Balvanera, P., Biggs, R., Cheung, W.W.L., Chini, L., Cooper, H.D., Gilman, E.L., Guénette, S., Hurtt, G.C., Huntington, H.P., Mace, G.M., Oberdorff, T., Revenga, C., Rodrigues, P., Scholes, R.J., Sumaila, U.R. & Walpole, M. (2010) Scenarios for Global Biodiversity in the 21st Century. Science, 330, 1496–1501.

Reside, A.E., VanDerWal, J., Phillips, B., Shoo, L., Rosauer, D., Anderson, B.A., Welbergen, J., Moritz, C., Ferrier, S., Harwood, T.D., Williams, K.J., Mackey, B., Hugh, S. & Williams, S.E. (2013a) Climate Change refugia for terrestrial biodiversity: the role of refugia ecosystem resilience and maintenance of terrestrial biodiversity in the face of global climate change. National Climate Change Adaptation Research Facility, Griffith University, Gold Coast, Qld.

Reside, A.E., VanDerWal, J., Phillips, B., Shoo, L.P., Rosauer, D., Anderson, B.J., Welbergen, J., Moritz, C., Ferrier, S., Harwood, T.D., Williams, K.J., Mackey, B., Hugh, S. & Williams, S.E. (2013b) Climate change refugia for terrestrial biodiversity. James Cook University.

Rosauer, D.F., Ferrier, S., Williams, K.J., Manion, G., Keogh, J.S. & Laffan, S.W. (2014) Phylogenetic generalised dissimilarity modelling: a new approach to analysing and predicting spatial turnover in the phylogenetic composition of communities. Ecography, 37, 21–32.

Sayre, R., Dangermond, J., Frye, C., Vaughan, R., Aniello, P., Breyer, S.P., Cribbs, D., Hopkins, D., Nauman, R., Derrenbacher, W., Wright, D.J., Brown, C., Convis, C., Smith, J.H., Benson, L., VanSistine, D.P., Warner, H., Cress, J.J., Danielson, J.J., Hamann, S.L., Cecere, T., Reddy, A.D., Burton, D., Grosse, A., True, D., Metzger, M., Hartmann, J., Moosdorf, N., Durr, H., Paganini, M., Defourny, P., Arino, O., Maynard, S., Anderson, M. & Comer, P. (2014) A new map of global ecological land units—An ecophysiographic stratification approach. Association of American Geographers.

Serrano, J., Richardson, J.E., Pennington, T.D., Cortes-B, R., Cardenas, D., Elliott, A. & Jimenez, I. (2018) Biotic homogeneity of putative biogeographic units in the Neotropics: A test with Sapotaceae. Diversity and Distributions, 24, 1121–1135.

Tittensor, D.P., Walpole, M., Hill, S.L.L., Boyce, D.G., Britten, G.L., Burgess, N.D., Butchart, S.H.M., Leadley, P.W., Regan, E.C., Alkemade, R., Baumung, R., Bellard, C., Bouwman, L., Bowles-Newark, N.J., Chenery, A.M., Cheung, W.W.L., Christensen, V., Cooper, H.D., Crowther, A.R., Dixon, M.J.R., Galli, A., Gaveau, V., Gregory, R.D., Gutierrez, N.L., Hirsch, T.L., Höft, R., Januchowski-Hartley, S.R., Karmann, M., Krug, C.B., Leverington, F.J., Loh, J., Lojenga, R.K., Malsch, K., Marques, A., Morgan, D.H.W., Mumby, P.J., Newbold, T., Noonan-Mooney, K., Pagad, S.N., Parks, B.C., Pereira, H.M., Robertson, T., Rondinini, C., Santini, L., Scharlemann, J.P.W., Schindler, S., Sumaila, U.R., Teh, L.S.L., van Kolck, J., Visconti, P. & Ye, Y. (2014) A mid-term analysis of progress toward international biodiversity targets. Science, 346, 241–244.

UN (2015) Transforming our world: the 2030 agenda for sustainable development. In: (ed. U. Nations), New York.

UNEP & WCMC (2016) Protected planet: Teh World Database on Protected Areas. In. UNEP-WCMC, Cambridge, UK.

Ware, C., Williams, K.J., Harding, J., Hawkins, B., Harwood, T., Manion, G., Perkins, G.C. & Ferrier, S. (2018) Improving biodiversity surrogates for conservation assessment: A test of methods and the value of targeted biological surveys. Diversity and Distributions, 24, 1333–1346.

Warren, D.L., Cardillo, M., Rosauer, D.F. & Bolnick, D.I. (2014) Mistaking geography for biology: inferring processes from species distributions. Trends in Ecology & Evolution, 29, 572–580.

Watson, J.E.M., Jones, K.R., Fuller, R.A., Marco, M.D., Segan, D.B., Butchart, S.H.M., Allan, J.R., McDonald-Madden, E. & Venter, O. (2016) Persistent Disparities between Recent Rates of Habitat Conversion and Protection and Implications for Future Global Conservation Targets. Conservation Letters, 9, 413–421.

Wilson, A.M. & Jetz, W. (2016) Remotely Sensed High-Resolution Global Cloud Dynamics for Predicting Ecosystem and Biodiversity Distributions. PLOS Biology, 14, e1002415.

