## Supplementary material for "Supporting global biodiversity assessment through high-resolution macroecological modelling: Methodological underpinnings of the BILBI framework": R code detailing analytical approach

run\_obspair\_GDM-Copy1


### **R Notebook for running an observation pair Generalised Dissimilarity Model**¶

#### This notebook accompanies the manuscript -¶

#### The intent of this notebook is to provide a worked example of the fitting procedure for an observation pair GDM model. Further detail on the model fitting proceedures can be found in the manuscript and supplementary documentation¶

##### Author: Andrew Hoskins.¶

#####¶

###### **Figure:** Example outputs from the BILBI moelling framework. This figure shows a visualisation of the spatial patterns of bet-diveristy on Mindinao Island in the Phillipines. Similar coloured areas represent areas of similar vertebrate communities. Overlayed are cleared areas (blacked out), protected areas (green polygons) and ecoregions (red lines)¶

In [1]:

```
## load supporting functions
source("./example/BILBI_supportingFunctions.R")
```

```
Loading required package: SDMTools
Warning message:
"package 'SDMTools' was built under R version 3.4.4"Loading required package: raster
Loading required package: sp

Attaching package: 'raster'

The following object is masked from 'package:SDMTools':

    distance
```

In [2]:

```
## load supporting packages
library(nnls)
library(SDMTools)
library(raster)
```

In [3]:

```
## load supporting data
load('./example/obsPair_GDM_example.RData')
## this will load a data.frame called Mod_data into the session. 
## This data.frame contains the random draw of species matches 
## and mismatahces and associated environmental covariates for 
## each site/pixel that the pair of species was drawn
```

In [4]:

```
## interrogate data
head(mod_data)
ls()
## Match - binary. was the pair of species a match or mismatch
## Lon1, Lat1, Lon2, Lat2 - Numeric. The geographic coordinates for each site
## For descriptions of the environmental variables refer to Table S2 in the supporting 
## documents of the manuscript.
## Richness.S1, Richness.S2 - integer. The number of unique speices found at each site. 
##                            While true richness at each site is unknown a filtered version
##                            of this is used as 'best-available' when assessing model fit
## SORENSON - numeric. Dissimilarity calculated as Bray-Curtis dissimilarity between sites
##            as with the richness calculations, this is not true but 'best-available' and
##            used only to show model fit in these examples
## TARGET - logical. True == site pair from within the target bio-realm, FALSE == site pair
##          from outside the target bio-realm. See S1.3 and main text for more detail
```

A data.frame: 6 × 39

|  | Match | Lon1 | Lat1 | Lon2 | Lat2 | EAAS | PTA | TNI | TRX | TXX | ... | BD\_2 | CLAY\_2 | PH\_2 | BARE\_2 | EPI\_2 | SILT\_2 | Richness.S1 | Richness.S2 | SORENSON | TARGET |
| --- | --- | --- | --- | --- | --- | --- | --- | --- | --- | --- | --- | --- | --- | --- | --- | --- | --- | --- | --- | --- | --- |
|  | <dbl> | <dbl> | <dbl> | <dbl> | <dbl> | <dbl> | <dbl> | <dbl> | <dbl> | <dbl> | ... | <dbl> | <dbl> | <dbl> | <dbl> | <dbl> | <dbl> | <dbl> | <dbl> | <dbl> | <lgl> |
| 1 | 1 | -71.57917 | 18.3208333 | -82.97083 | 9.729167 | 1127.113 | 1176 | 16.0 | 13.080553 | 31.90774 | ... | 1545.075 | 42.425 | 5.0650 | 1 | 109.65679 | 25.45 | 14 | 29 | 1 | TRUE |
| 3 | 1 | -92.59583 | 15.3541667 | -84.01250 | 10.437500 | 1418.062 | 3337 | 16.7 | 16.411995 | 35.20415 | ... | 1214.325 | 30.350 | 4.7450 | 1 | 106.48545 | 37.55 | 2 | 6 | 1 | TRUE |
| 4 | 1 | -78.49583 | 0.3541667 | -70.53750 | 19.245833 | 1276.974 | 1509 | 11.4 | 13.025219 | 24.42522 | ... | 1651.575 | 28.100 | 6.3175 | 1 | 83.20821 | 24.55 | 3 | 2 | 1 | TRUE |
| 7 | 1 | -93.17917 | 17.5875000 | -42.97917 | -22.429167 | 1376.002 | 3820 | 17.0 | 13.306551 | 34.70655 | ... | 1356.575 | 30.350 | 5.5375 | 1 | 54.12861 | 17.40 | 45 | 22 | 1 | TRUE |
| 8 | 1 | -88.67917 | 17.1541667 | -95.04583 | 17.320833 | 1392.381 | 2348 | 17.9 | 9.395161 | 30.90923 | ... | 1594.575 | 31.350 | 5.0325 | 1 | 83.52948 | 31.55 | 4 | 1 | 1 | TRUE |
| 9 | 1 | -88.99583 | 16.7958333 | -84.07083 | 10.070833 | 1338.923 | 2011 | 16.0 | 9.495810 | 29.60971 | ... | 913.400 | 21.500 | 5.5450 | 1 | 99.80428 | 42.55 | 1 | 2 | 1 | TRUE |

1. 'fitGDM'
2. 'I\_spline'
3. 'inv.logit'
4. 'mod\_data'
5. 'negexp'
6. 'nnls.fit'
7. 'nnnpls.fit'
8. 'obs.gdm.plot'
9. 'ObsTrans'
10. 'plotSpline'
11. 'splineData'

In [5]:

```
## The observation pair GDM follows standard GDM by using monotonic regression splines to describe the non-linear
## responses of ecological communities to environmental variation. Due to the spatially nested design of the
## analytical approach, with target biorealms sourrounded by neighbouring buffering regions we fit using either 
## 3, 4 or 5 splines to each modelling domain.
##
## 3 splines are used when all environmental envelope is covered withing the target modelling domain.
## 4 splines are used when the envionrmental envelope exceeds the range of the modelling domain in one direction
## 5 splines are used when the environmental envelope exceeds the range of the modelling domain in both directions

## fold data from both sites into single vectors representing the full range of observed environmental valeues
## Note: bare ground is a binary value, as such, it is not included in the splining process
p1 <- mod_data[,c(6:17,19:20)]
p2 <- mod_data[,c(21:32,34:35)]
colnames(p2) <- colnames(p1)
preds <- rbind(p1,p2)

## extract environmental values from target modelling domain and buffering regions
targetPreds <- preds[mod_data$TARGET,]
bufferPreds <- preds[!mod_data$TARGET,]

## calculate quantiles for each spline along 
q1 <- unlist(lapply(1:ncol(preds),function(x){min(bufferPreds[,x])}))
q5 <- unlist(lapply(1:ncol(preds),function(x){max(bufferPreds[,x])}))
q2 <- unlist(lapply(1:ncol(preds),function(x){min(targetPreds[,x])}))
q4 <- unlist(lapply(1:ncol(preds),function(x){max(targetPreds[,x])}))
q3 <- unlist(lapply(1:ncol(preds),function(x){quantile(targetPreds[,x],0.5)}))

## logical test. Do target and buffering quantiles match 
test1 <- q1 < q2
test5 <- q4 < q5

## using logical test create vectors indicating the number of splines used (3, 4 or 5) per environmental variable,
## and the position of each spline.
quans <- c()
spl <- c()
for(i in 1:length(q1)){
    if(test1[i] & test5[i]){q <- c(q1[i],q2[i],q3[i],q4[i],q5[i]);s <- 5}
	if(!test1[i] & !test5[i]){q <- c(q1[i],q3[i],q5[i]);s <- 3}
	if(test1[i] & !test5[i]){q <- c(q1[i],q2[i],q3[i],q5[i]);s <- 4}
	if(!test1[i] & test5[i]){q <- c(q1[i],q3[i],q4[i],q5[i]);s <- 4}
	if(any(duplicated(q))){q <- q[!duplicated(q)]; s <- length(q)}
	quans <- c(quans,q)
	spl <- c(spl,s)
	}

## clean up redundant objects
rm(list=c("p1","p2","preds","q1","q2","q3","q4","q5"))
gc()
```

A matrix: 2 × 6 of type dbl

|  | used | (Mb) | gc trigger | (Mb) | max used | (Mb) |
| --- | --- | --- | --- | --- | --- | --- |
| Ncells | 5184611 | 276.9 | 8273852 | 441.9 | 5539395 | 295.9 |
| Vcells | 148653886 | 1134.2 | 279172008 | 2130.0 | 250711830 | 1912.8 |

In [8]:

```
## Now that we know the position of each spline it's time to transform the environmnetal data using a monotonic
## regression splines fitted at the positions calculated in the previous steps. This uses the support function splineData
## This function takes three arguements;
## data - data.frame or matrix. contains the data to be splined
## splines - vector - the number of splines top be fitted to each variable
## quantiles - vector - the position of each spline

## create data.frame containing only the environmental variables
toSpline <-  mod_data[,c(6:17,19:20,21:32,34:35)]

## spline data
splinedData <- splineData(toSpline,splines=spl,quantiles=quans)

## check outputs
head(splinedData)
```

A matrix: 6 × 59 of type dbl

| EAAS\_spl1 | EAAS\_spl2 | EAAS\_spl3 | EAAS\_spl4 | PTA\_spl1 | PTA\_spl2 | PTA\_spl3 | PTA\_spl4 | TNI\_spl1 | TNI\_spl2 | ... | PH\_spl4 | EPI\_spl1 | EPI\_spl2 | EPI\_spl3 | EPI\_spl4 | SILT\_spl1 | SILT\_spl2 | SILT\_spl3 | SILT\_spl4 | SILT\_spl5 |
| --- | --- | --- | --- | --- | --- | --- | --- | --- | --- | --- | --- | --- | --- | --- | --- | --- | --- | --- | --- | --- |
| 0 | 2.117088e-02 | 0.40638796 | 0.197945482 | 0 | 0.23069698 | 0.27246080 | 0.004929266 | 0 | 0.0009835550 | ... | 0 | 0 | 0.0160453302 | 0.397838687 | 0.14853958 | 0 | 5.981317e-03 | 0.02179082 | 0.000000000 | 0 |
| 0 | 0.000000e+00 | 0.07018553 | 0.151920130 | 0 | 0.00000000 | 0.10925886 | 0.038802096 | 0 | 0.0000885195 | ... | 0 | 0 | 0.0000000000 | 0.110951101 | 0.23161426 | 0 | 7.181853e-03 | 0.36380079 | 0.085832540 | 0 |
| 0 | 5.266509e-03 | 0.08379259 | 0.000000000 | 0 | 0.09603913 | 0.03513286 | 0.000000000 | 0 | 0.0308442889 | ... | 0 | 0 | 0.0261454231 | 0.423236359 | 0.12230420 | 0 | 1.589748e-02 | 0.39975268 | 0.089317788 | 0 |
| 0 | 9.194189e-03 | 0.23586498 | 0.032566026 | 0 | 0.04506075 | 0.42006947 | 0.058851035 | 0 | 0.0583149728 | ... | 0 | 0 | 0.2810304065 | 0.324851005 | 0.00000000 | 0 | 1.777988e-01 | 0.52268768 | 0.054794887 | 0 |
| 0 | 4.285220e-03 | 0.20963092 | 0.047969393 | 0 | 0.00000000 | 0.03519417 | 0.001855732 | 0 | 0.0002458887 | ... | 0 | 0 | 0.0009721999 | 0.003685228 | 0.00000000 | 0 | 0.000000e+00 | 0.05702609 | 0.009519364 | 0 |
| 0 | 6.296181e-05 | 0.06265171 | 0.008681271 | 0 | 0.00516956 | 0.17803096 | 0.009678585 | 0 | 0.0393028528 | ... | 0 | 0 | 0.0257245264 | 0.295157426 | 0.01550446 | 0 | 8.651421e-05 | 0.41337295 | 0.193770752 | 0 |

In [9]:

```
## prepare data for model fitting
## data stricture consists of the binary response variable and all environmental
## predictor variables. All predictors are splined with the exception of bare ground
data <- as.data.frame(cbind(Match=mod_data$Match,splinedData))
data$BARE <- abs(mod_data$BARE - mod_data$BARE_2) ## prepare bare ground variable

## create formula to pass to model fitting function
f1 <- paste(colnames(data)[-1],collapse="+")
f1_formula <- as.formula(paste(colnames(data)[1],"~",f1,sep=""))
f1_formula
```

```
Match ~ EAAS_spl1 + EAAS_spl2 + EAAS_spl3 + EAAS_spl4 + PTA_spl1 + 
    PTA_spl2 + PTA_spl3 + PTA_spl4 + TNI_spl1 + TNI_spl2 + TNI_spl3 + 
    TNI_spl4 + TNI_spl5 + TRX_spl1 + TRX_spl2 + TRX_spl3 + TRX_spl4 + 
    TXX_spl1 + TXX_spl2 + TXX_spl3 + TXX_spl4 + TXX_spl5 + WDI_spl1 + 
    WDI_spl2 + WDI_spl3 + WDI_spl4 + WDX_spl1 + WDX_spl2 + WDX_spl3 + 
    WDX_spl4 + TWI_SAGE_spl1 + TWI_SAGE_spl2 + TWI_SAGE_spl3 + 
    TWI_SAGE_spl4 + RUG_spl1 + RUG_spl2 + RUG_spl3 + BD_spl1 + 
    BD_spl2 + BD_spl3 + BD_spl4 + CLAY_spl1 + CLAY_spl2 + CLAY_spl3 + 
    CLAY_spl4 + CLAY_spl5 + PH_spl1 + PH_spl2 + PH_spl3 + PH_spl4 + 
    EPI_spl1 + EPI_spl2 + EPI_spl3 + EPI_spl4 + SILT_spl1 + SILT_spl2 + 
    SILT_spl3 + SILT_spl4 + SILT_spl5 + BARE
```

In [10]:

```
## fit model - stage 1
## Stage 1 identified coeffcents for all splined environmental variables within the model
## model fitting uses a logit link and binomial error distribution
## following traditional GDM analyses, positive coefficients are forced for all predictor
## variables. The exception in an observation pair GDM is that the intercept is allowed
## to be negative, allowing for the model to fit in logit space which ranges both positive
## and negative rather than the traditional negative exponential link function whose space
## is only positive.
fit_1 <- fitGDM(formula=f1_formula,data=data,family=binomial(),method='nnnpls.fit')

## inspect model
summary(fit_1)
## Note: because the form of the model only allows positive coefficients - ensuring a positive
## relationship between ecological distance and environmental distance - there will be no
## negative coefficients and where a positive relationship was not found, the returned
## coefficient will be zero.

## extract coefficients
coefs_1 <- coefficients(fit_1)
```

```
Call:
glm(formula = formula, family = family, data = data, control = list(maxit = 500), 
    method = method)

Deviance Residuals: 
    Min       1Q   Median       3Q      Max  
-3.2533  -0.9866  -0.2016   1.0330   2.1206  

Coefficients:
               Estimate Std. Error  z value Pr(>|z|)    
(Intercept)   -2.163757   0.004978 -434.651   <2e-16 ***
EAAS_spl1      0.000000  51.184001    0.000    1.000    
EAAS_spl2      0.000000   0.025184    0.000    1.000    
EAAS_spl3      0.757969   0.014053   53.936   <2e-16 ***
EAAS_spl4      0.000000   0.026897    0.000    1.000    
PTA_spl1       0.000000  51.183926    0.000    1.000    
PTA_spl2       0.000000   0.026923    0.000    1.000    
PTA_spl3       0.000000   0.031628    0.000    1.000    
PTA_spl4       0.000000   0.135811    0.000    1.000    
TNI_spl1       0.000000   0.360421    0.000    1.000    
TNI_spl2       0.000000   0.039254    0.000    1.000    
TNI_spl3       1.304736   0.014310   91.179   <2e-16 ***
TNI_spl4       0.000000   0.017050    0.000    1.000    
TNI_spl5       0.000000   1.077476    0.000    1.000    
TRX_spl1       0.298627   0.021452   13.921   <2e-16 ***
TRX_spl2       0.686166   0.015190   45.173   <2e-16 ***
TRX_spl3       0.000000   0.034783    0.000    1.000    
TRX_spl4       0.000000   0.231006    0.000    1.000    
TXX_spl1       0.000000   0.300638    0.000    1.000    
TXX_spl2       0.000000   0.037527    0.000    1.000    
TXX_spl3       0.948901   0.013089   72.494   <2e-16 ***
TXX_spl4       0.000000   0.014450    0.000    1.000    
TXX_spl5       0.000000   0.055093    0.000    1.000    
WDI_spl1       0.000000   0.036039    0.000    1.000    
WDI_spl2       0.358981   0.012641   28.398   <2e-16 ***
WDI_spl3       1.772836   0.022477   78.875   <2e-16 ***
WDI_spl4       0.000000   0.097267    0.000    1.000    
WDX_spl1       0.000000   0.085533    0.000    1.000    
WDX_spl2       0.271638   0.019689   13.796   <2e-16 ***
WDX_spl3       0.261281   0.022729   11.496   <2e-16 ***
WDX_spl4       0.000000   0.070781    0.000    1.000    
TWI_SAGE_spl1  0.000000   0.015630    0.000    1.000    
TWI_SAGE_spl2  0.249990   0.012081   20.693   <2e-16 ***
TWI_SAGE_spl3  0.000000   0.018307    0.000    1.000    
TWI_SAGE_spl4  0.000000   0.145361    0.000    1.000    
RUG_spl1       0.259645   0.006903   37.615   <2e-16 ***
RUG_spl2       0.000000   0.037096    0.000    1.000    
RUG_spl3       0.000000   0.275827    0.000    1.000    
BD_spl1        0.000000   0.030707    0.000    1.000    
BD_spl2        0.398783   0.014856   26.843   <2e-16 ***
BD_spl3        0.000000   0.022059    0.000    1.000    
BD_spl4        0.000000   0.272942    0.000    1.000    
CLAY_spl1      0.000000   0.459529    0.000    1.000    
CLAY_spl2      0.000000   0.026004    0.000    1.000    
CLAY_spl3      0.505562   0.020391   24.793   <2e-16 ***
CLAY_spl4      0.000000   0.096458    0.000    1.000    
CLAY_spl5      0.307321   0.440301    0.698    0.485    
PH_spl1        0.000000   0.069394    0.000    1.000    
PH_spl2        0.604800   0.019616   30.832   <2e-16 ***
PH_spl3        0.000000   0.021696    0.000    1.000    
PH_spl4        0.000000   0.204397    0.000    1.000    
EPI_spl1       0.000000   0.123577    0.000    1.000    
EPI_spl2       0.000000   0.017756    0.000    1.000    
EPI_spl3       1.452421   0.011085  131.030   <2e-16 ***
EPI_spl4       0.000000   0.016645    0.000    1.000    
SILT_spl1      0.000000   1.561373    0.000    1.000    
SILT_spl2      0.400809   0.021630   18.530   <2e-16 ***
SILT_spl3      0.617453   0.016971   36.384   <2e-16 ***
SILT_spl4      0.000000   0.036422    0.000    1.000    
SILT_spl5      0.113522   0.375309    0.302    0.762    
BARE           0.000000   0.011451    0.000    1.000    
---
Signif. codes:  0 '***' 0.001 '**' 0.01 '*' 0.05 '.' 0.1 ' ' 1

(Dispersion parameter for binomial family taken to be 1)

    Null deviance: 2772589  on 1999999  degrees of freedom
Residual deviance: 2403559  on 1999939  degrees of freedom
AIC: 2403681

Number of Fisher Scoring iterations: 4
```

In [11]:

```
## fit model - stage 2
## Stage 2 fits to the relationship between geographic distance and the residual unexplained
## variation from the stage 1 fitting procedure. In this case, a linear function is fitted
## however non-linear functions could also be used during this stage.

## calculate distance between the species observations in each observation pair.
dist <- SDMTools::distance(mod_data$Lat1,mod_data$Lon1,mod_data$Lat2,mod_data$Lon2)
## get the linear predictor from the stage 1 model fitting
eco.intRemM <- fit_1$linear.predictors
## setup data
dataTab <- data.frame(Match=fit_1$y,Ecological=eco.intRemM,Distance=dist$distance/1000)
## fit model - linear predictor from stage 1 is used as an offset in the fitting of stage 2
## ensuring that the model can only interact with the residual variation from stage 1
fit_2 <- glm(Match ~ offset(Ecological) + Distance,family=binomial(),data=dataTab,control=list(maxit=1000),method='nnls.fit')

## inspect model
summary(fit_2)

## extract coefficients
coefs_2 <- coefficients(fit_2)
```

```
Call:
glm(formula = Match ~ offset(Ecological) + Distance, family = binomial(), 
    data = dataTab, control = list(maxit = 1000), method = "nnls.fit")

Deviance Residuals: 
    Min       1Q   Median       3Q      Max  
-3.4633  -1.0503  -0.2109   0.8671   2.1203  

Coefficients:
             Estimate Std. Error z value Pr(>|z|)    
(Intercept) 0.000e+00  2.331e-03     0.0        1    
Distance    1.726e-04  1.068e-06   161.6   <2e-16 ***
---
Signif. codes:  0 '***' 0.001 '**' 0.01 '*' 0.05 '.' 0.1 ' ' 1

(Dispersion parameter for binomial family taken to be 1)

    Null deviance: 2403559  on 1999999  degrees of freedom
Residual deviance: 2342406  on 1999998  degrees of freedom
AIC: 2342410

Number of Fisher Scoring iterations: 4
```

In [12]:

```
## inspect the fitted functions from the observation pair GDM.

## setup plotting region
par(mfrow=c(2,2))

## find the position of each coefficient 
pos <- c(0,cumsum(spl))

## plot for each of the 14 environmental predictor variables when a relationship was found
for(n in 1:14){
    nSplines <- spl[n]
    pCoeffs <- coefs_1[(pos[n]+2):(pos[n+1]+1)]
    pQuants <- quans[(pos[n]+1):pos[n+1]]
    env <- seq(pQuants[1],pQuants[nSplines],length.out=100)
    xlab <- gsub("_spl1","",names(pCoeffs[1])) ## fix xlab here - why pCoeffs 1 not always 1
    ylab <- paste("f(",xlab,")")
    if(any(pCoeffs != 0)){plotSpline(env,nSplines,pCoeffs,pQuants,xlab,ylab)}    
}

## plot of the relationship between species observation matches and geographic distance
rng_dist <- range(dist$distance/1000)
xx <- seq(rng_dist[1],rng_dist[2],length.out=100)
yy <- xx*coefs_2[2]
yy_link <- inv.logit(coefs_1[1]+coefs_2[1] + yy)
par(mfrow=c(2,1))
plot(xx,yy,xlab="Geographic distance between sites (km)",ylab="Ecological distance",type="l")
plot(xx,yy_link,xlab="Geographic distance between sites (km)",ylab="Linked ecological distance",type="l")
```

In [13]:

```
## inspect model fit. 
source("./example/BILBI_supportingFunctions.R")
obs.gdm.plot(fit_2,"Fitted observation pair GDM",w=440.117673210066,Is=coefs_1[1]+coefs_2[1],raw=mod_data)

## This function generates a 4 panel plot giving the relationship between the observed binary response
## variable and the resulting fitted values, densities of the species matches (0) and species mismatches (1)
## along the fitted environmental gradient, and an estimate of the relationship between communitiy disimilarity
## and the the predicted dissimilarity - however because true dissimilarity is unknown this is estimated
## from pre-calculated values using a basic filter (each site must have > 15 unique species).
## The top left panel shows the observed proportion of species missmatches (including binomial variance) across
## 10 bins vs the predicted proportion of species missmatches. 
## Top right is the density of observed species matches (red) and missmatches (blue) across the gradient of 
## predicted ecological distance - we expect a well fitting model to show some seperation between these two 
## densities.
## Bottom left is the observed (blue) and predicted (green) proportion of species missmatch along the predicted 
## ecological gradient.
## Bottom right shows the estimated fit of community dissimilarity vs the proportion of species missmatches for
## observed (blue) and predicted (green) values
```

In [18]:

```
##------------------------------------------------------------------------------------------------------------------------##
##------------------------------------------------------------------------------------------------------------------------##
##------------------------------------------------------------------------------------------------------------------------##
##------------------------------------------------------------------------------------------------------------------------##
##------------------------------------------------------------------------------------------------------------------------##
```

### Example analytical process - Calculate biodiversity persistence and extinction risk.¶

#### In this example we will use the results from the previously fitted observation pairs GDM to estimate biodivesrity persistence for a given grid cell - i.e. the proportion of species within a grid cell that is expected to ersist anywhere within their range.¶

#### For an an example of the BILBI framework being used to estimate persistence globally see:¶

> ## \*Di Marco M., Harwood T.D., Hoskins A.J., Ware C., Hill S.L.L, Ferrier S. (2019). Projecting impacts fo global climate and land-use scenarios on plant biodiveisty using compositional-turnover modelling. Global Change Biology. DOI: 10.1111gcb.14663\*

#### Note: The BILBI modelling framework uses highly optimised C++ code, custom data structures and CSIRO's high performance computing infrastructure to rapidly perform these analyses for every cell across the terrestrial surface of the globe. This workable example is coded in R to enable users to run the analysis on a moderately powerful desktop computer. Users attempting to run a global scale analysis with this code should budget sufficient time for the anlaysis to run.¶

In [19]:

```
## load transformed grid data structure. These are grids consisting of a non-linear transform 
## of each environmental grid to include the estimated coefficients. See paper below for further
## details.
##    Manion G., (2009) A technique for monotonic regression splines to enable non-linear 
##       transformation of GIS rasters. 18th World IMACS/MODSIM Congress, Cairns, Australia.
##       https://www.mssanz.org.au/modsim09/F13/manion_F13a.pdf
trans_env <- brick("./example/Example_transgirds_crop.grd")
```

In [20]:

```
## load habitat condition layer. This is a combination of the estimated impact of land-use to 
## local biodiversity (via the PREDICTS project) and a downscaled version of the land-use
## harmonisation v2 dataset. For more details on these datasets and methods see
##
##    Purvis A., Newbold T., De Palma A., Contu S., Hill S.L.L., Sanchez-Ortiz K., Phillips H.R.P.,
##       Hudson L.N., Lysenko I., Borger L., Scharlemann J.P.W. (2018). Modelling and projecting the
##       response of local terrestrial biodiversity worldwide to land use and related pressures: the
##       PREDICTS project. Advances in Ecological Research. Vol 58. pp 201-241.
##
##    Hoskins AJ., Bush A., Gilmore J., Harwood T.D., Hudson L.N., Ware C., Williams., Ferrier S.
##       (2016) Downscaling land-use data to provide global 30" estimates of five land-use classes.
##       Ecology and Evolution. Vol 6. Issue 9.
##
##    Di Marco M., Harwood T.D., Hoskins A.J., Ware C., Hill S.L.L, Ferrier S. (2019).
##       Projecting impacts of global climate and land-use scenarios on plant biodiveisty
##       using compositional-turnover modelling. Global Change Biology. 
##       DOI: 10.1111gcb.14663
cond <- raster("./example/Example_HabitatCondition_crop.grd")
```

#### Calculating biodiversity persistence and extinction risk.¶

#### Biodiversity persistence is an estimate of the proportion of the species associated with a cell that are expected to persits anywhere within their natural range. It's inverse is extinction risk which is an extimate of the proportion of species associated with a cell that are committed to extinction within their natural range.¶

##### Biodiversity persistence is calculated for a single cell using the following equation:¶

$$p\_i=\Bigg[\frac{\sum\_{j=1}^n s\_i,\_j c\_j}{\sum\_{j=1}^n s\_i,\_j}\Bigg]^z$$

##### where:¶

$s\_i,\_j =$ the modelled similarity in species composition from an observation pair GDM,

$c\_j =$ the estimated condiction of habitat condition within each of those cells,

##### and¶

$z =$ the coefficient of the species area relationship

##### Extinction risk is the inverse and is calculated as follows:¶

$$e\_i=1-p\_i$$

###### Note: To allow for an example that will function on most desktop machines, this example only works with an approximately 5 deg x 5 deg area of the full biorealm. This means that while the estimates will approximate results from BILBI they will not match them exactly.¶

In [24]:

```
## extract required variables

z <- 0.25 ## assumed species area relationship following Di Marco et al (2019) and references contained within.

## extract the intercept from the full model
intercept <- coefs_1[1] + coefs_2[1]

## geographic distance coefficiets
g_dist_coef <- coefs_2[2]

## how many cells in the raster layer
cells <- length(cond)

## how many comparison cells
n_comp <- 100000
```

In [25]:

```
## because without the aid of high performance computing we cannot run this across all cells, this example shows
## the calculation run for a single focal cell that is compared against 100,000 random cells from the region.

s_i <- sample(cells,1) ## focal cell
s_j <- sample(cells,n_comp) ## comparison cells
s_j <- s_j[s_j != s_i] ## make sure we're not comparing with the same cell

## samples from focal cell
xy_i <- xyFromCell(cond,s_i)
env_i <- extract(trans_env,xy_i)

## samples from comparison cells
xy_j <- xyFromCell(cond,s_j)
env_j <- extract(trans_env,xy_j)
tf <- !is.na(env_j[,1])
env_j <- env_j[tf,]
cond_j <- extract(cond,xy_j[tf,])
```

In [26]:

```
## calculate environmental distance between focal cell and sample cells
env_i_exp <- matrix(ncol=nrow(env_j),nrow=ncol(env_j))
env_i_exp[1:length(env_i_exp)] <- env_i
env_i_exp <- t(env_i_exp)

env_dist <- rowSums(abs(env_j - env_i_exp))

## calculate geographic distance component
dist_ij <- SDMTools::distance(rep(xy_i[2],nrow(env_j)),rep(xy_i[1],nrow(env_j)),xy_j[tf,2],xy_j[tf,1])$dist / 1000
dist_ij_coef <- g_dist_coef * dist_ij

## build full linear predictor
lin_pred <- intercept + dist_ij_coef + env_dist ## 

## link out of logit space
pred <- inv.logit(lin_pred)

## transform into dissimilarity
diss <- ObsTrans(p0=inv.logit(intercept),w=440.117673210066,p=pred)$out
```

In [33]:

```
## calculate persistence
persistence <- (sum((1-diss)*cond_j) / sum(1-diss))^z

cat("Persistence: ",persistence,"\n")

## calculate extinction risk
extinction <- 1- persistence

cat("Extinction risk: ",extinction)
```

```
Persistence:  0.9596196 
Extinction risk:  0.04038039
```
